# Decoding frontotemporal and cell type-specific vulnerabilities to neuropsychiatric disorders and psychoactive drugs

**DOI:** 10.1101/2022.11.30.518630

**Authors:** Jiatong Ji, Honglu Chao, Huimei Chen, Jun Liao, Yangfan Ye, Yongping You, Ning Liu, Jing Ji, Enrico Petretto

## Abstract

Abnormalities in temporal and frontal lobes (TL and FL) have been linked to cognition and neuropsychiatric disorders. While structural and functional differences between the brain lobes have been documented in disease, the cellular heterogeneity in FL and TL and its impact to the vulnerability to genetic risk factors for neuropsychiatric disorders is not well studied. We hypothesize that intrinsic cellular-level differences between TL and FL explain the vulnerability of specific cell types to genetic risk factors and psychoactive drugs. To test this, we integrated single-nucleus transcriptome analysis in fresh human FL and TL with data related to genetic susceptibility and gene dysregulation in neuropsychiatric disease, and response to psychoactive drugs. We also investigate how these differences are associated with gene dysregulation in disease brain. Neuronal cell populations were the most vulnerable to psychiatric genetic risk factors, and more specifically parvalbumin interneurons (PVALB neurons). These PVALB-expressed genetic risk factors were mostly upregulated in the TL compared with FL, and dysregulated in the brain of patients with obsessive-compulsive disorder, bipolar disorder and schizophrenia. We found *GRIN2A* and *HCN1*, implicated in schizophrenia by genome-wide association studies, to be significantly upregulated in PVLAB from the TL and in brain cortex from schizophrenia patients. Our analysis provides comprehensive evidence for PVALB neurons as the most vulnerable cell type that is implicated in several psychiatric disorders. PVALB neurons showed the highest vulnerability to psychoactive drug response, which was 3.6-fold higher than the vulnerability to genetic risk factors. In summary, we show high vulnerability of PVALB neurons that is specific to the temporal lobe, implying that differences between TL and FL greatly influence the cell vulnerability to genetic risk factors as well as the response to psychoactive drugs. These findings offer insights into how regional brain differences affect the cell type vulnerabilities in neuropsychiatric disorders.

## INTRODUCTION

Genome-wide association studies (GWAS) elucidated the genetic etiology and pathophysiology of neuropsychiatric disorders^1^, identifying a multitude of associated loci and disease risk factors (aka genetic susceptibility factors), and revealing substantial genetic complexity^2, 3^. Several clinical studies have shown that diverse neuropsychiatric disorders share similar symptoms, which, at least in part, could be due to common genetic factors underlying these disorders^4^, which impact central functional processes such as neuronal regeneration, myelination, synapse maturation and plasticity^5, 6^.

In addition to the role played by genetics, the FL and TL are believed to be involved in the pathophysiology of neuropsychiatric disorders, including bipolar disorder (BP), schizophrenia (SCHI), and ASD^7-9^. For example, changes in cortical volume, surface area, and thickness have been reported in in schizophrenia patients, who show significant reduction of the entire inferior temporal gyrus, while bipolar patients showed significant reduction of cortical thickness in main TL as well as part of FL and parietal lobe^10^. Schizophrenic patients also exhibit functional, structural, and metabolic abnormalities in the prefrontal cortex (PFC)^11^. Other studies showed structural alterations in the hippocampus and in medial TL regions in patients with schizophrenia, bipolar disorder, and depression^12^. However, most of these studies focused on structural brain changes between FL and TL in patients with psychiatric disorders, and a link between these structural changes and dysregulation of specific cellular processes and/or cell vulnerabilities in disease have not firmly established yet.

The regional differences between TL and FL might also affect gene expression in specific cell types, and in turn these might impact important functional processes dysregulated in neuropsychiatric disorders. For example, region-specific characteristics of neurons in the cortex, including different electrophysiological features, have been previously reported^13^, and region-specific transcriptomic changes in neurons from the cerebral cortex (and the subcortical region) showed specific patterns of molecular and developmental regulation^13^. Cellular level differences between brain regions might be also relevant to disease pathobiology; e.g., excitatory and inhibitory neurons in the middle frontal gyrus showed a significant association with neuropsychiatric disorders^14^, and further cell type and brain area associations have been reported for depressive disorder^15^. In autism spectrum disorders (ASD) patients, the expression level of synaptic and neurodevelopmental genes in layer 2/3 cortical neurons from PFC is especially affected^16^. However, given the very high cellular complexity within and between brain regions such as TL and FL, cell type-specific and regional-specific gene expression changes remain to be fully elucidated.

Single cell/nuclei sequencing approaches now permit to analyze and “chart” disease-associated gene expression changes at the cellular level. For example, specific cell types associated with schizophrenia and anorexia nervosa have been identified by integrating mouse snRNA-seq of brain with GWAS data^17^. However, the relationship between the vulnerable cell type in psychiatric disorders and the cellular processes involved, especially the frontotemporal cortex, is less well studied and remain poorly understood. Previous studies integrated snRNA-seq analysis with existing drug-specific signatures to determine cell type-specific vulnerabilities^18^, but the relative contribution of genetic risk factors and psychoactive drugs to cell type vulnerability, and the effect for brain regional differences, have not been characterized. To bridge this knowledge gap, here we integrated single-nucleus RNA sequencing (snRNA-seq) data from fresh brain tissue samples of TL and FL, with (*i*) GWAS data for 7 psychiatric disorders, (*ii*) RNA-seq data from healthy and disease human brain cortex and (*iii*) drug-specific gene targets for common psychoactive drugs. This allowed us to define a detailed map of single cell changes associated with brain regional differences (TL and FL), and assess how these affect cell type vulnerability to both psychiatric genetic risk factors and psychoactive drugs. We also linked changes in TL and FL cell types to gene dysregulation in human brain tissue from patients with neuropsychiatric disorders. The integrated datasets and results presented here provide a unique resource to help understanding the cell-specific changes during disease pathogenesis and pharmacological treatments using common psychoactive drugs.

## METHODS

### Tissue collection and library preparation

Three human brain samples were obtained from patients without any psychiatric disorder history in the First Affiliated Hospital of Nanjing Medical University, which was approved by the Research Ethics Committee of Nanjing Medical University (Nanjing, Jiangsu, China), and experiments were performed in accordance with the approved guidelines. The detail information and the sampling site are in Table S3 and Figure S10 respectively. Three normal tissues were obtained from healthy part during surgery of multiple intracranial small space-occupying diseases. When selecting sampling site, we used preoperative MRI images to select a gray matter of the cerebral cortex that was far away from the primary lesion and edema. Liquid nitrogen is used to inhibit the tissue from degradation. All the experiments were performed in accordance with the approved guidelines.

Single-nucleus RNA-sequencing (snRNA-seq) analysis was performed by NovelBio Co.,Ltd. The tissues were surgically removed and snap frozen in liquid nitrogen for intact nucleus isolation. The nucleus isolation was carried out by Nuclei Isolation Kit: Nuclei EZ Prep (Sigma, NUC101). Briefly, the frozen tissue was homogenized in NLB buffer which contain 250 mM Sucrose, 10 mM Tris-HCl, 3 mM MgAc2, 0.1% Triton X-100 (SigmaAldrich, USA), 0.1 mM EDTA, 0.2U/μL RNase Inhibitor (Takara, Japan). Various concentration of sucrose was used to purify the nucleus. The concentration of nucleus was adjusted to about 1000 nuclei/μL for snRNA-Seq. The snRNA-Seq libraries were generated using the 10X Genomics Chromium Controller Instrument and Chromium Single Cell 3’V3.1 Reagent Kits (10X Genomics, Pleasanton, USA). Briefly, cells nuclei were concentrated to 1000 nuclei/μL then loaded into each channel to generate single-cell Gel Bead-In-Emulsions (GEMs). After the RT step, GEMs were broken and barcode-cDNA was purified and amplified. The amplified barcoded cDNA was fragmented, A-tailed, ligated with adaptors and index PCR amplified. The final libraries were quantified using the Qubit High Sensitivity DNA assay (Thermo Fisher Scientific, USA) and the size distribution of the libraries were determined using a High Sensitivity DNA chip on a Bioanalyzer 2200 (Agilent, USA). All libraries were sequenced by Novaseq6000 (Illumina, USA) on a 150 bp paired-end run.

### Preprocessing of single cell RNA sequencing data

Raw reads were processed using the CellRanger software (v.1.3.1). The “cellranger count” command was used to generate the gene-per-cell expression matrices for each sample by aligning the reads to the genome and quantifying expression of the reference genes (GRCh38). Three single-cell gene expression matrices corresponding to the three biological samples were generated. SnRNA-seq data were mainly processed by R package Seurat (v4.0.2)^19^. The expression matrices of the 3 samples were merged into a global Seurat object using the “MergeSeurat” function. The gene expression profiles of each single cell were then merged together yielding 56,082 cells. Then coming to the quality control routine, a cut-off value of 300 and 4,000 unique molecular identifiers (UMIs) were used to select single cells for further analysis. Cells with potential mitochondrial content >10% were removed. Next cells outside the 5th and 95th percentile of the distribution of the number of detected genes were discarded. Then we filtered out doublets, which were identified using DoubletFinder^20^. Finally, we obtained 45,229 cells. The data were normalized for library size by a scale factor of 10,000 and log-transformed using the ‘NormalizeData’ function in Seurat, followed by ‘ScaleData’. Highly variable features were determined using the function ‘FindVariableFeatures’ from Seurat. Principal component analysis (PCA) was first performed to obtain a small number of principal components as input to the UMAP algorithm. The number of principal components was then chosen at the ‘elbow’ of the plot where a substantial drop was observed in the proportion of variance explained.

### Batch correction and UMAP plot generation

In order to remove any potential batch effects between TL and FL samples, we ran the CCA analysis^19^. First, we split the dataset into 2 groups according to the brain region before eliminating the batch effect. Functions used for this step were ‘FindIntegrationAnchors’ and ‘IntegrateData’ from Seurat. Then, we ran ‘ScaleData’, ‘RunPCA’, ‘FindNeighbors’, ‘RunUMAP’ and ‘FindClusters’. To determine the optimal cluster resolution, we first clustered the data on a range of resolutions (from 0.1 to 2, with steps of 0.2). Then, we plotted the number of obtained clusters versus the resolution and chose a resolution (i.e., 0.3) for which the number of clusters remained constant across multiple resolutions. For visualization, we ran ‘DimPlot’ in Seurat to generate UMAP plot.

### Cell type identification and KEGG pathways annotation

Differential gene expression analyses between cell clusters were performed using logistic regressions on the integrated normalized counts, by the ‘FindAllMarkers’ function in Seurat with the following settings: assay = ‘RNA’, only.pos=TRUE, test.use = ‘LR’, min.pct = 0.1, logfc.threshold = 0.25. This procedure was used for subcluster classification. Each cluster was then aggregated into main cell type groups by comparing the top 20 genes in each cell types with the known markers^21^. To corroborate the cell classification, we also used the BRETIGEA package, which provides a well-validated set of brain cell type-specific marker genes derived from multiple types of experiments, including marker specificity, enrichment, and absolute expression^22^. A hypergeometric test was used to test the overlap between our data-driven markers and the top cell type markers from BRETIGEA. To identify the subcluster-specific genes within excitatory neuron (EX) and inhibitory neuron (INH), differential gene expression analysis between subclusters was performed. We assigned EX subclusters to respective cortical layers by evaluating the expression of layer-specific markers identified by^23^. INH formed two major types (branches), associated with different developmental origins in caudal ganglionic eminence (CGE) and middle ganglionic eminence (MGE), respectively^24^. The MGE included PVALB, SST subclasses and the CGE contained LAMP5, VIP subclasses^25^. To interpret the differential gene expression results we carried out KEGG pathway enrichment. To annotate the specific functions of each cell type or neuron-subcluster, we ranked all the DEGs between clusters detected above by avg_log_2_FC in descending order and then input them into ‘gseKEGG’ function of the clusterProfiler^26^ package in R. To remove redundancy in these pathway annotations, the significant pathways (BH-adjusted P <0.05) were then merged and clustered using the default clustering method in pheatmap package.

### Differential expression analysis in FL vs TL and pathway enrichment analysis

For each group of cells, we extracted counts from the processed snRNA-seq data. In order to remove the bias from unbalance proportion of the 2 regions, we did down sampling for each cell type to make the cell number consistent across brain regions. Then we performed differential expression using edgeR (v3.20.8) to identify TF/FL region-associated genes (threshold of absolute log fold change>0.5 and false discovery rate (FDR)<0.05). In the analysis, ‘calcNormFactors’ was used to adjust for varying sequencing depths (due by differing library sizes) as well as minimize the log-fold changes between the samples for most genes. We the used ‘estimateDisp’ function to estimate common dispersion and tagwise dispersions and lastly, the grouped data (FL vs TL) were input to ‘glmQLFTest’ function. After obtaining all the DEGs in each cell type, we combined them together as a matrix and performed hierarchical clustering of the FC(TL/FL) and further annotated the genes function by functional enrichment analysis by ‘enrichKEGG’ in clusterProfiler (threshold for significance: adjusted P <0.05)^26^. Combining the results of the hierarchical clustering with the functional enrichment, we identified six clusters of DEGs that show (1) cell type-specific gene expression patterns between FL and TL and (2) a coherent biological function (reported in Figure 2c). The thresholds used for finding the DEGs and the enriched pathways in each neuron-subcluster are the same as above analysis.

### Enrichment of neuropsychiatric disease GWAS genes in brain cell clusters

To identify whether the neuropsychiatric disease genes identified by GWAS show cell type specific association or regional alterations between FL and TL, we downloaded a list of GWAS candidate genes related to bipolar disorder (BP), autism spectrum disorder (ASD), schizophrenia (SCHI), ADHD, obsessive compulsive disorder (OCD), unipolar depression (UD), cognitive impairment (CI) from the NHGRI-EBI GWAS catalog^27^. We only considered the GWAS genes that have been detected by published studies that pass the NHGRI-EBI GWAS catalogue eligibility criteria. Since one GWAS SNP variant can be “tagged” to multiple genes within the same associated locus, we chose to focus on the genes reported in at least one published study as well as detected (expressed) in our brain snRNA-seq data. After filtering out the redundant genes, we selected gene associations with P-value ≤ 10^−5^. A summary of the number of GWAS genes selected in each disease is shown Figure 1a.

**Fig. 1.**
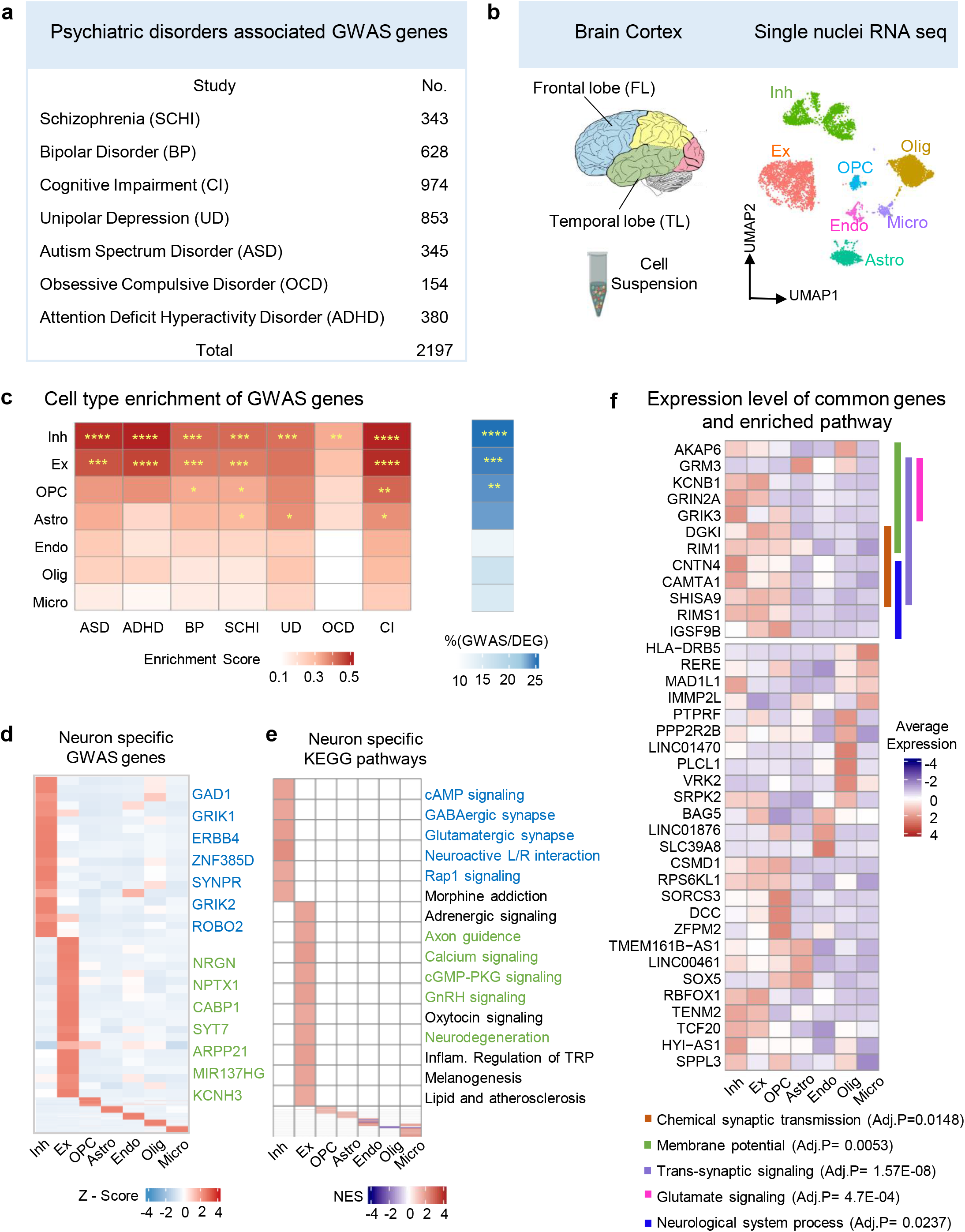
GWAS-associated genes for neuropsychiatric disorders are enriched for neuronal expression. **a**. Number of GWAS genes for neuropsychiatric disorders extracted from NHGRI-EBI GWAS catalogue. **b**. Schematic diagram displaying the TL and FL brain neocortex (*left*). UMAP plot showing all main cell types identified in the snRNA-seq analysis: astrocyte (Astro) oligodendrocyte (Olig), microglia cells (Micro), endothelial cells (Endo), oligodendrocyte progenitor cells (OPC), excitatory neuron (EX) and inhibitory neuron (INH)(*right*). More details in Figure S1. **c**. Cell type enrichment score for expression of neuropsychiatric-associated GWAS genes grouped by disease (*left*). Proportion of GWAS genes within marker genes of each type with significance (BH-adjusted P-value) of the hypergeometric test labelled by asterisks (*right*): Significance (P-value) thresholds: * 0.01-0.05, ** 0.001-0.01, *** 0.0001-0.001, **** <0.0001. **d**. Heatmap showing the specific marker genes that are also GWAS-associated genes in EX (green) and INH (blue), respectively. **e**. Gene set enrichment analysis (GSEA) of cell type-specific differential expression in EX and INH (BH-adjusted P<0.05). Pathways in green and in blue are contributed by GWAS genes in EX and INH, respectively. **f**. Average (normalized) expression level of 38 genes (detected in 7 neuropsychiatric diseases) in each cell type; the significant (non-redundant) GO biological processes (BH-adjusted P<0.05) for these 38 genes are also indicated by different colors (vertical bars) and annotated at the bottom.

Two enrichment tests were performed. (1) To test for the enrichment in gene expression of GWAS genes in cell types, we calculated the average expression levels of each gene set (e.g., GWAS genes for a given disease) and derived the module score for these GWAS genes in each cell type by the ‘Addmodulescore’ function in Seurat^19^. The module score was transformed into z-score, and the p-value of the z-score was calculated by the “pnorm” function in R. (2) To test the enrichment (overrepresentation) of GWAS genes in each cell type, we used a hypergeometric test for the overlap between cell type specific genes and the GWAS genes in each disease. In both cases, the background gene set used in the hypergeometric test was all genes expressed in each cell type (or sub-type), and the P-values of enrichment were corrected for multiple testing using the Benjamini-Hochberg (BH) procedure.

### Functional network of GWAS genes for neuropsychiatric diseases

After filtering the GWAS genes, we explored the enriched biological pathways. We carried out KEGG pathway enrichment analysis by ‘enrichKEGG’ function of the clusterProfiler^26^ package in R. The significant pathways (BH-adjusted P<0.05) were then grouped into multiple pathway clusters according to their high-level pathway name entry in the KEGG database. Many genes are shared by different pathways and this property was assessed by means of Jaccard similarity coefficient. For each pathway pair, the Jaccard similarity coefficient was calculated by assessing the number of intersect genes between 2 pathways divided by the number of union genes in 2 pathways. The pathways’ relationships are represented as a network, where each edge connecting any two pathways is proportional to the Jaccard similarity coefficient; the resulting network was plotted by Cytoscape software (v 3.9.0).

### Comparison with genes dysregulated in disease brain tissue

To identify genes dysregulated in the brain tissue from patients with neuropsychiatric disease, we retrieved public datasets from the Gene Expression Omnibus (GEO) database, as well as retrieving preprocessed lists of DEGs from original publications. For the GEO data, we downloaded three bulk-RNA datasets related to: MDD (GSE102556)^28^, ASD (GSE64018)^29^, BP (GSE 53239)^30^. For OCD and SCHI, we downloaded the list of DEGs from ^31^ and ^32^, respectively. “edgeR” was used to perform differential expression analysis by comparing gene expression in control group vs disease group. After correction of multiple testing, genes with BH-adjusted P-value<0.05 and absolute log_2_FC>0.25 were selected as disease associated genes. For each disorder, the list of DEGs (disease associated genes) was test for overlap with set of DEGs between FL and TL identified here. The significance of the overlap was assessed by hypergeometric test (background: all genes expressed in each cell type or sub-type) and the P-values were corrected using the BH approach, as indicated.

### Gene co-expression networks of GWAS genes enriched for expression in neurons

For selected subset of GWAS-genes enriched in neuronal subtypes we derive cell type specific gene co-expression networks (Figure 2g, Figure 2i, Figure 4f). We used Spearman correlation to calculate the connection between each gene, where the correlations in gene expression were calculated across cells in the same compartments (e.g., within inhibitory neurons). We used the Spearman correlation as this performs better than traditional Pearson correlation and has with higher accuracy and reproducibility for network reconstruction using single cell data^33^. All the gene-gene connections with P<0.05 for the Spearman correlation were deemed significant and retained to represent the gene network, which was plotted by Cytoscape software (v 3.9.0). In the network graph, each edge connecting any two genes represents the Spearman correlation.

**Fig. 2.**
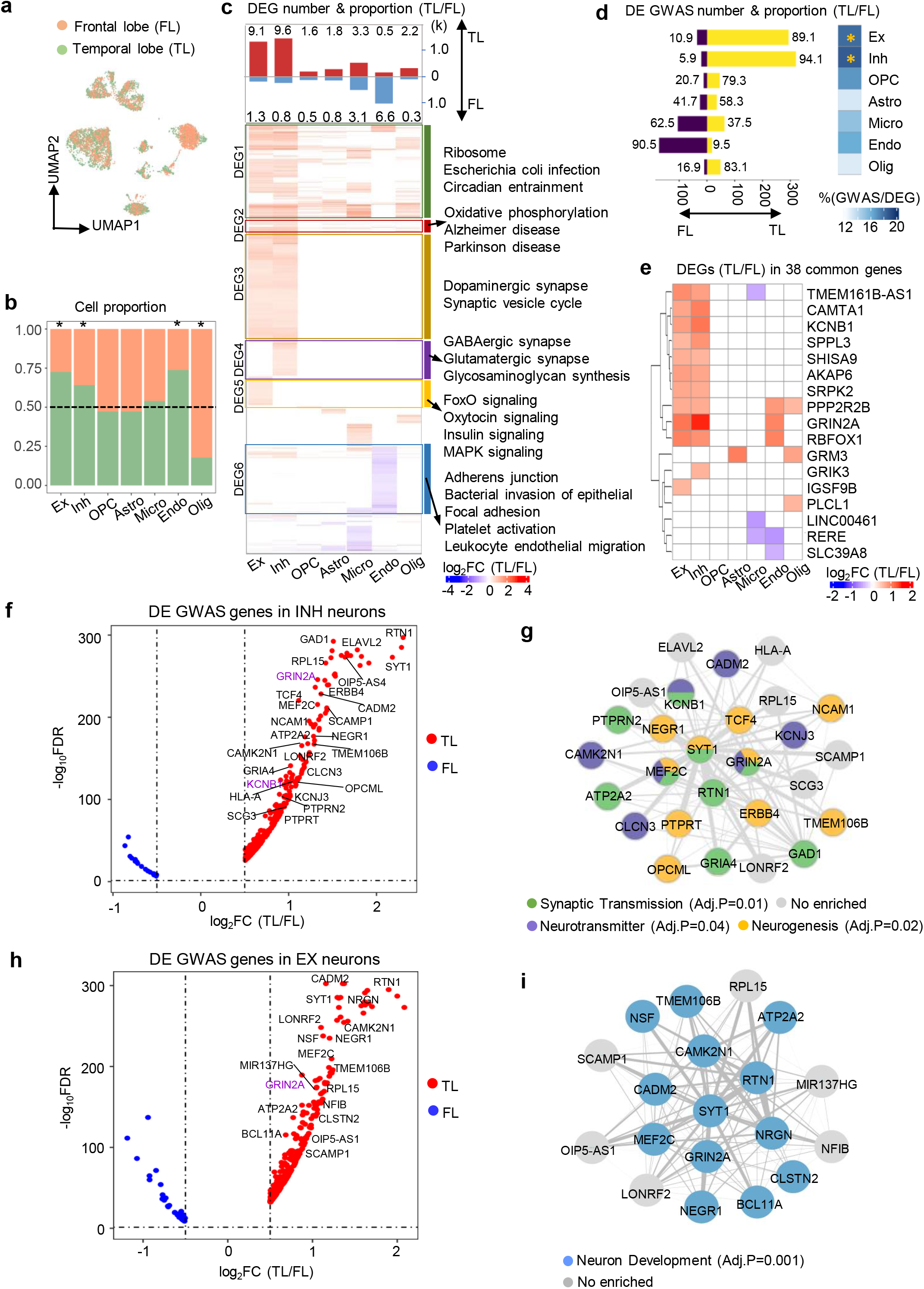
GWAS-associated genes for neuropsychiatric disorders are enriched for neuronal expression in the temporal lobe. **a**. UMAP plot showing how the two regions overlap after batch correction (see Methods). **b**. Bar plot showing the cell type proportions in TL and FL samples. Asterisks mark significantly different proportions by chi-square test (P<0.05). **c**. Number of DEGs (absolute log_2_FC >0.5 and FDR<0.05) between FL and TL, split by up regulated in TL and FL, respectively. For each cell type, the proportion (%) of DEGs to the total number of expressed genes is indicated (*top*). Hierarchical clustering of log fold change of DEGs between FL and TL identified in each cell type, which are grouped into 6 clusters according to the KEGG pathways annotation (BH-adjusted P<0.05) (*bottom*). **d**. Number of GWAS genes that are DE between FL and TL, split into up-regulated in TL (yellow) and in FL (purple), respectively. Numbers represent the ratio (%) of the up-regulated DE GWAS genes in each region over the total DE GWAS genes (*left*). Proportion of all DE GWAS genes over the total number of DEGs in each cell type, and the significance of enrichment in each cell type by hypergeometric test (*right*). *, BH-adjusted P<0.05 **e**. Heatmap showing the log_2_FC(TL/FL) of DEGs (absolute log_2_FC>0.5 and FDR<0.05) between FL and TL among 38 GWAS genes (detected in 7 neuropsychiatric diseases). **f, h**. Volcano plots displaying GWAS gene expression in EX (h) and INH (f) neurons, respectively. Genes with log_2_FC>1 and GWAS P<10^−8^ are highlighted. Genes with color highlight belong to the 38 common genes detected in 7 disorders. **g, i**. Gene networks describing the interaction of genes which are highlighted in the corresponding volcano plots. Edges represent gene-gene co-expression in the specific cell type. Each node (gene) in the network is colored according to the significant GO functional enrichment (BH-adjusted P<0.05).

### Enrichment of psychoactive drug target genes in brain cell types

In order to evaluate the response to psychotropic medications in each brain cell type, drug target gene sets were downloaded from^18^, originally retrieved from DSigDB (http://dsigdb.tanlab.org/DSigDBv1.0/). Eight drug classes, which have been approved by clinical trials and are widely used in the clinic (with good efficacy in the treatment of psychiatric disorders), have been considered here: SSRI (Selective Serotonin Reuptake Inhibitor), DA (Dopamine Receptor Agonists), TCA (Tricyclic Antidepressant), BZDs (Benzodiazepines), MAOI (Monoamine Oxidase Inhibitor), GRA (Glutamate Receptor Agonists), GABA (GABA Receptor Agonists), and ARA (Adrenergic receptor agonist). With respect to depression and bipolar disorder, we included classical antidepressants (SSRI, TCA, MAOI, BZDs), and they also contribute to schizophrenia treatment. ARA is used in bipolar, autism and depression treatment. With respect to autism, and ADHD, OCD, GRA, and GABA are commonly used. Although DA is mainly used for Parkinson patients, many studies indicate its efficiency in BP and schizophrenia too^34, 35^. The enrichment of drug target genes was performed in the same way used to assess enrichment of GWAS genes (detailed above). The background gene set used in the enrichment analysis was all genes expressed in each cell type or sub-type, and the P-values were corrected using the BH approach, as indicated.

## RESULTS

### Susceptibility genes for neuropsychiatric disorders share common pathways

To investigate which functional and cellular processes are dysregulated in neuropsychiatric disorders, we collected GWAS data from 7 psychiatric disorders (BP, ASD, SCHI, attention deficit hyperactivity disorder (ADHD), obsessive compulsive disorder (OCD), unipolar depression (UD), and cognitive impairment (CI)), which have been directly or indirectly linked to processes dysregulated in TL and/or FL (Figure 1a and Table S1). A subset of 38 genes was detected by GWAS in all 7 neuropsychiatric disorders (Figure S1a), and these were enriched for KEGG pathways and biological processes, including synapse related as well as neuron degeneration diseases, endocrine function and neuronal signaling pathways (Figure S1b). We also annotated a set of “core pathways” shared between at least 4 neuropsychiatric diseases (Figure S1c, Table S2). Amongst the most common pathways, synaptic dysfunction and neurodevelopmental conditions (*blue circle* in Figure S1c) have been reported to be altered in postmortem brain in ASD^36^. Besides, exome sequencing studies suggested the role of post-synaptic density in SCHI^37, 38^. Many signaling transduction associated pathways, including AKT, AMP and AMPK, play a role in higher mental functions such as mood, neurogenesis and neurotransmitter^39, 40^ (*green circle* in Figure S1c). We also found that GnRH-related pathways belonging to endocrine system being enriched in most of the disorders (*pink circle* in Figure S1c); hormones regulate ovary and testicular function in mammals and changes in gonadotropin levels are known to impact mood and contribute to the development of affective disorders^41^. These analyses show that GWAS genes for heterogenous neuropsychiatric diseases impact commonly shared pathways and broad brain functions important for disease pathobiology. To shed light on potential regional and cell type specific functions we carried out single cell profiling in the brain.

### Susceptibility genes for neuropsychiatric disorders are enriched for expression in neuronal cell types

We sequenced single nuclei from frozen brain cortex tissues of 3 subjects (one sample from the dorsolateral prefrontal cortex in FL, and two samples from temporopolar lobes) with no history of neuropsychiatric disorder (Figure 1b, Table S3), and, after filtering out poorly sequenced nuclei and potential doublets, we identified a total of 45,229 cells (Figure S2a). Uniform manifold approximation and projection (UMAP) identified 7 main cell clusters, which were also manually corroborated using well-known marker genes (Figure S2b). Excitatory neurons were the most abundant (38.3%) cell type, followed by inhibitory neurons and oligodendrocyte (23.6% and 22.1%, respectively), which is consistent with previously reported cell proportions in cortical tissue areas^42^ (Figure S2c-d). Excitatory neurons were characterized by the expression of *ENC1* and *SLC17A6*^43^, while inhibitory neurons were identified by GABAergic (*GAD2, GRIK1*) and dopaminergic neurons by *TH*^43-45^. Immune cells displayed a specific expression of well-known markers, e.g., *CD74* in microglia^46^, *RGS5* in endothelial cells^47^. *PTPRZ1* and *BCAN* were highly expressed in OPC^48, 49^, while oligodendrocytes are characterized by *MOBP* and *MBP* ^50^ expression. Expression of *AQP4* was characteristic for astrocytes^51^. Pathway enrichment analysis confirmed functional specialization of different cell types (Figure S2e), supporting the fidelity of our brain cell types clustering.

Next, we integrated the neuropsychiatric disorders GWAS data with brain snRNA-seq data to annotate the cell type specific expression and cellular mechanism of the disease susceptibility genes, and detail whether these genes exhibit expression differences between FL and TL. Overall, all genes associated with disease by GWAS (thereafter named “GWAS genes”) (n=2,197, combined across 7 diseases) showed the highest enrichment for expression in EX, INH and OPC (adjusted P-value = 8×10^−9^, 10^−4^, 3×10^−3^, respectively, hypergeometric test, Figure 1c). GWAS genes accounted for 21.7% and 24.6% of the neuronal marker genes in EX and INH, respectively. Considering each disease separately, we found significant enrichment for all diseases in INH markers, and for 5 diseases in EX markers (Figure 1c). We also identify GWAS genes such as *NRGN, CABP1, NPTX1* as cell type markers for EX, and *GRIK1, GAD1, ERBB4* for INH (Figure 1d). These results show that amongst other brain cells, a significantly large proportion of GWAS genes are expressed in neurons, suggesting this cell type might play a key role in mediating the genetic susceptibility to several neuropsychiatric disorders^52^.

The pathways impacted by the GWAS genes expressed specifically in EX and INH neurons are consistent with the 4 common pathway groups enriched in the whole set of GWAS genes (“core pathways” in Figure S1b and S1d), which is in keeping with previous reports^53^. The main pathways are axon guidance, calcium signaling process, GnRH signaling pathway, and neuro-degeneration diseases for the EX-expressed GWAS genes, and GABAergic synapse, neuroactive L/R interaction and depression for the INH-expressed GWAS genes (Figure 1e). When we looked at the set of 38 GWAS genes shared across all 7 psychiatric disorders, not surprisingly, we found a significant enrichment for expression only in EX and INH neurons (adjusted P-value of 0.004 and 6.4×10^−5^, respectively, hypergeometric test). Furthermore, a subset of 12 genes (most highly expressed in EX and INH neurons) shows specific and non-redundant enrichment for Gene Ontology (GO) biological processes. Specifically, *GRIN2A, KCNB1*, GRM3, *GRIK3* are enriched in synaptic signaling and glutamate signaling; *DGK1, RIM1* are related to synaptic transmission, *SHISA9* and *RIMS1* contribute to neural processes, while *CNTN4* and *CAMTA1* are involved in both (Figure S1c and S1f).

### Extensive cell type-specific transcriptional changes in brain cortex between frontal and temporal lobes

We then investigated the cell types and expression changes between FL and TL (Figure 2a). EX, INH and Endo showed the highest cell proportion in TL, while Olig were overrepresented in FL (Figure 2b). We found the highest number of differentially expressed genes (DEGs) between FL and TL in EX and INH (>1,800 in each cell type), with most genes being upregulated in TL (Figure 2c, *top panel*). The larger number of DEG in TL vs FL in INH and EX was not affected by the higher proportion of INH and EX neurons in TL (see Methods). We also identified six clusters of DEGs (Figure S3) that show (1) coherent expression changes between FL and TL and (2) were enriched for the same biological functions or pathways (Figure 2c, *bottom panel*). These patterns reflect coherent changes in expression of genes, which were mostly upregulated in TL, especially in EX and INH.

Resolving these regional gene expression changes at the single-cell level might help elucidating the functional context of GWAS genes for neuropsychiatric disorders. We therefore investigated whether the GWAS genes overlap with the DEGs between FL and TL identified in each cell type. The DEGs in EX and INH showed the largest overlap with GWAS genes (adjusted P-value <0.05, hypergeometric test) and most of them are upregulated in TL compared to FL (Figure 2d, *bar plot*). Of the 38 common GWAS genes, 17 (45%) showed significant changes between TL and FL, with most genes (>89%) being upregulated in TL in EX and INH, including *KCNB1, GRIN2A, CAMTA1* (Figure 2e), and enriched for biological processes such as regulation of neurological system, glutamate signaling and synaptic signaling (Figure S4h).

We then explored the relationship between the significance of GWAS associations (GWAS P-value) and the degree of change (fold change, FC) in gene expression between TL and FL in EX and INH. While we found no obvious and consistent correlations between GWAS gene significance and FCs (Figure S4a-g), we identified *GRIN2A* and *KCNB1* with highly significant GWAS associations (P<10^−8^) in all 7 diseases, and more strongly upregulated TL (FDR<0.05, log_2_FC(TL/FL)>1) in either both INH/EX (*GRIN2A*) or in INH (*KCNB1*) (Figure 2f, h). The cell type-specific gene co-expression networks describing the interactions between the most significant GWAS genes (P<10^−8^) were enriched for neurogenesis and neurotransmitter in INH (Figure 2g), and neuron development in EX (Figure 2i). Therefore, we showed a significantly high proportion of genetic risk factors for neuropsychiatric disorders are upregulated in neuronal cells from the TL.

### Susceptibility genes for neuropsychiatric disorders are mostly enriched in PVALB interneurons from the temporal lobe

Given the results above we focused on neuronal cells. Using our single cell transcriptomic data to identify neuronal cell sub-types, we uncovered EX-subtypes L2-3, L4, L5, L6, and INH-subtypes SST, PVALB, LAMP5, VIP (Figure 3a-b). Neuron subtypes were identified by well-established markers that also enriched for specific functional categories^24^ (Figure S5 a-b). To identify the most relevant neuronal sub-cluster for a particular disorder, we calculated the percentage of the GWAS genes overlapping with the cell type sub-cluster specific markers (Figure 3c). Overall, we found stronger enrichment in INH than in EX sub-clusters, with INH PVALB, INH SST and EX L5 being the neuronal sub-clusters mostly enriched for expression of GWAS genes, which accounted for about 33% of the marker genes in each sub cell type (Figure 3c, *right panels*). The overlap between GWAS genes and neuronal sub-cluster marker genes is also found at the pathway level. We found biological processes and pathways enriched in the marker genes of the neuronal sub-clusters (Figure 3d-e; *green highlight*) which are consistent with those identified in the GWAS genes (Figure S1b). For example, INH PVALB and SST were both enriched for processes related to neuron system development, synaptic neurotransmitter and glutamatergic signaling, which are also significantly enriched in GWAS genes (Figure S1b).

**Fig. 3.**
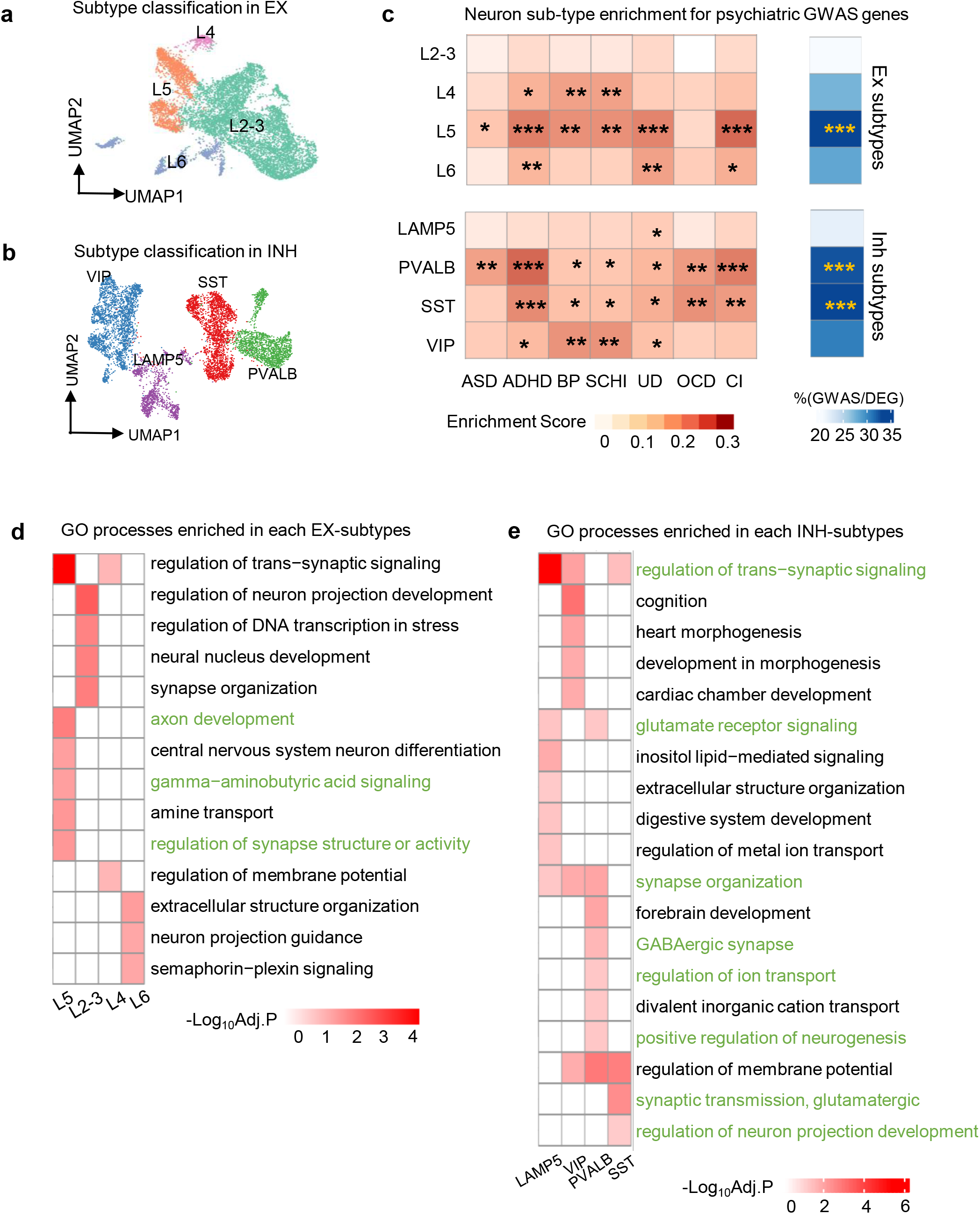
Specific neuronal subtypes show enrichment for GWAS-associated genes for neuropsychiatric disorders. **a, b**. UMAP plot showing the subtypes of EX (a) and INH (b) neurons. EX neurons are classified into L2-3, L4, L5, L6. INH neurons are classified into LAMP5, PVALB, SST, VIP. **c**. Cell type enrichment score for expression of neuropsychiatric disorder associated GWAS genes. Proportion (%) of GWAS genes within marker genes of each cell-subtype alongside significance of enrichment (adjusted P-value) by hypergeometric test (*right*). Significance (P-value) thresholds: * 0.01-0.05, ** 0.001-0.01, *** 0.0001-0.001, **** <0.0001 **d, e**. Enriched GO biological processes (BH-adjusted P<0.05) in each subtype of EX (d) and INH (e) neurons. Pathways highlighted in green match the pathways identified in GWAS genes (reported in Figure S1 and Table S2).

We went on to investigate the contribution of TL and FL differences on neuronal sub-clusters. For INH, cells from 2 regions are well mixed and we found no difference in cell number (Figure 4 a-b). Differential expression analysis between TL and FL in INH sub-clusters identified manifold DEGs, with the PVALB and LAMP5 sub-clusters having the highest number of DEGs (Figure 4c, *bar plot on the top panel*). Similarly, we characterized regional expression differences in EX neurons (Figure S6a-b), and detected many DEGs in EX sub-clusters, e.g., L5 and L4 (Figure S6c, *bar plot on the top panel)*. For both neuronal cell types, most of the DEGs were upregulated in TL compared with FL. Functional analysis of the marker genes in each INH sub-cluster shows enrichment for both common (e.g., “ribosome”) and sub-cluster specific processes (e.g., synaptic vesicle cycle in PVALB) (Figure 4c, *bottom panel*). Amongst other neuronal cells, the most specific functional enrichment was detected for PVALB cub-cluster, including signaling factor like GABA, calcium, cAMP, PKG that are essential for structural changes in synapses, dendrites and axons (Figure 4c, *bottom panel*), which are consistent with previous reports on structural plasticity differences between FL and TL in PVALB neurons^54^. Long-term potentiation and synaptic vesicle which can impact synapse and axon growth also differ in PVALB cluster^55^. These gene expression changes in functional pathways between TL and FL would be consistent with those reported in neuropsychiatric diseases, for example, changes in long-term potentiation can induce depression, bipolar and schizophrenia^56^, degradation of cAMP or cGMP can result in depressive-like behaviors^57^.

**Fig. 4.**
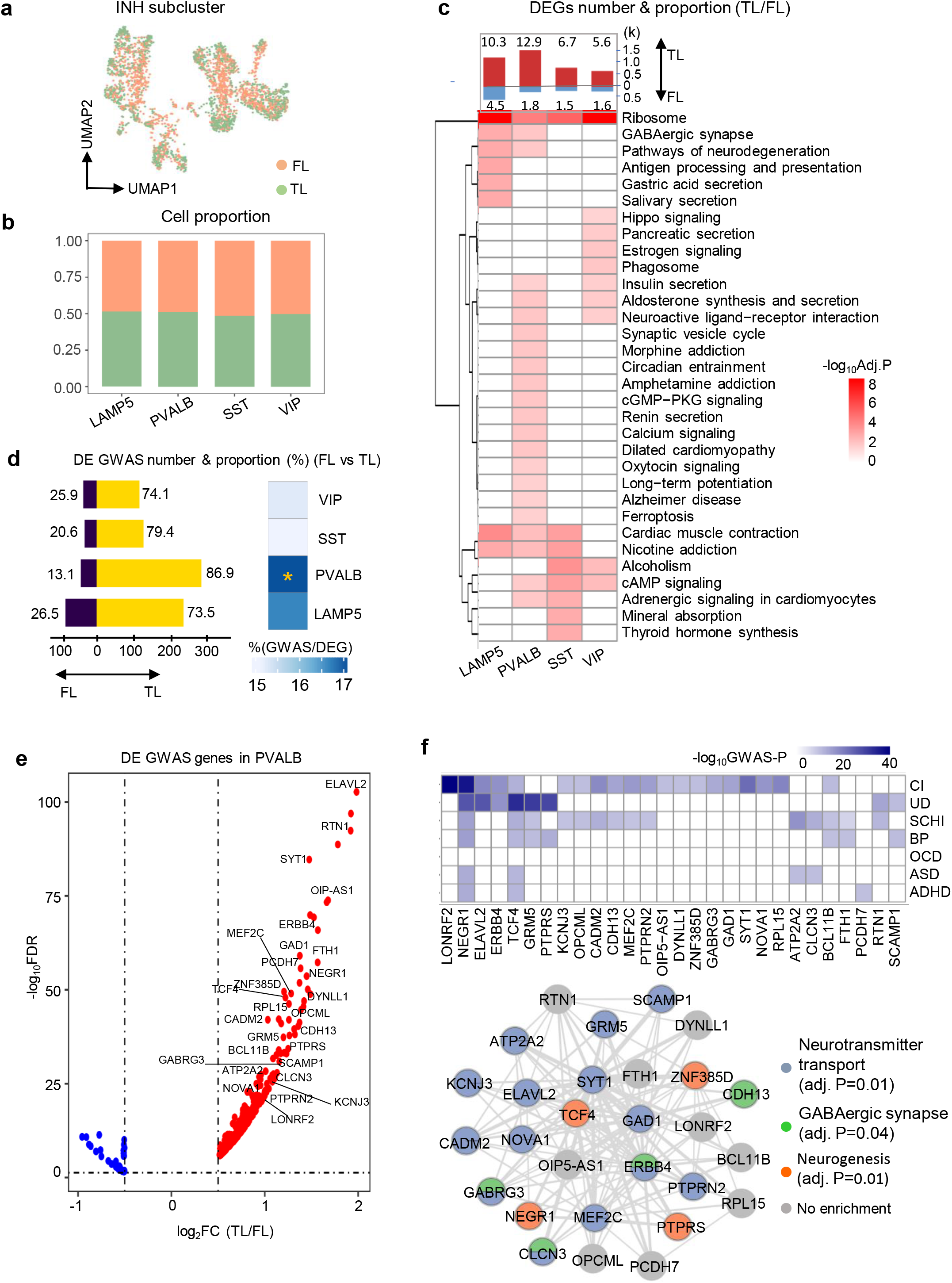
GWAS-associated genes for neuropsychiatric disorders are specifically enriched for PVALB expression in the temporal lobe. **a**. UMAP plot showing the FL and TL distribution to INH neurons. **b**. Cell proportion of FL and TL in each subcluster. **c**. Number of DEGs between FL and TL (absolute log_2_FC>0.5 and FDR<0.05) split by genes upregulated in FL and TL, respectively. The proportion (%) of DEGs over the total number of expressed genes is indicated for each subcluster (*top*). Heatmap exhibit the KEGG pathways enriched by DEGs between Fl and TL (BH-adjusted P<0.05). **d**. Number of GWAS genes that are DE between FL and TL, split by upregulated in TL (yellow) and FL (purple), respectively. Numbers represent the proportion (%) of the upregulated DE GWAS genes in each region over the total number DE GWAS genes (*left*). Proportion (%) of all DE GWAS genes over the total of DEGs in each cell type; significant enrichment (by hypergeometric test) is indicated (*right*). *, BH-adjusted P<0.05. **e**. Volcano plot showing log_2_FC(TL/FL) for the GWAS-associated genes in PVALB cluster. Subset of 28 GWAS genes with log_2_FC>1 and GWAS P<10^−8^ are highlighted in the graph. **f**. Heatmap showing the GWAS P-values for the 28 genes in each disease (*top*). Gene network describing the interaction of the 28 GWAS genes. Edges represent gene-gene co-expression in PVALB neurons. Each node (gene) in the network is colored according to the significant GO functional enrichment (BH-adjusted P<0.05).

To investigate whether gene expression differences in FL and TL at the neuronal sub-cluster level can influence neuropsychiatric disorders genes activity, we looked at the GWAS genes that are DE between TL and FL, and test for enrichment in each neuronal sub-cluster. While no enrichment was detected in any EX sub-cluster (Figure S6d), we found a significant enrichment of GWAS DEGs (n=327, ∼17%; enrichment P<0.05) in INH PVALB sub-cluster, with most GWAS genes (∼87%) being up regulated in TL (Figure 4d, e). Specifically, SCHI, BP and UD disorders had the highest enrichment for GWAS genes that are up-regulated in TL compared with FL (Figure S7a), including a subset of 11 GWAS genes that are more strongly expressed in PVALB neurons (Figure S7b). A set of 28 genes expressed in PVALB sub-cluster had robust significance of GWAS association (i.e., P<10^−8^) and sizeable difference in gene expression between TL and FL (i.e., log_2_FC>1) (Figure 4e). Overall, we found 28 distinct genes (out of 285 GWAS genes, 9.8%) that were upregulated in PVALB from the TL and robustly associated (GWAS P<10^−8^) with psychiatric disease (Figure 4f, *top panel*). We annotated the GWAS genes’ functional connectivity in PVALB cells and derived their PVALB-specific co-expression network (Figure 4f, *bottom panel*), which is enriched for neurotransmitter, neurogenesis, and GABAergic synapse processes. Dysregulation of these processes has been linked to the pathogenesis of mental diseases, e.g., PV-deficient GABA synapses increased asynchronous release of GABA and shape behavioral responses, which decreased the level of excitation, as observed in schizophrenia and related psychiatric disorders^58^.

### Psychoactive drugs target genes are enriched in PVALB interneurons from the temporal lobe

Additionally, we investigated the transcriptional impact (aka “vulnerability”^18^) of common and approved by clinical trials psychoactive drugs (SSRI: Selective Serotonin Reuptake Inhibitor, DA: Dopamine Receptor Agonists, TCA: Tricyclic Antidepressant, BZDs: Benzodiazepines, MAOI: Monoamine Oxidase Inhibitor, GRA: Glutamate Receptor Agonists, GABA: GABA Receptor Agonists, ARA: Adrenergic receptor agonist) on genes expressed in in specific cell types and sub cell types from the TL and FL^18^. To this aim, we used the drug-target dataset from Smail et al^18^, and assessed the degree to which the gene targets of each drug are expressed in specific cells and sub-cell types (see Methods). The drugs and gene targets for each psychoactive drug class are listed in Table S4. Drug target genes for SSRI and TCA were strongly overrepresented in the set of PVALB genes upregulated in TL (Table S4), and enriched for high expression (BH-adjusted P<0.05) in PVALB cells specifically from the TL (Figure S8a-b). Overall, we report 42 (out of 118, 35.6%) genes upregulated in PVALB from TL that are responsive to many of the psychoactive drugs considered here (Table S4). We also found a significant enrichment for drug targets genes in other neuronal subtypes. However, compared with other neuronal sub-types, the expression level of drug-target genes was higher in PVALB from TL and, differently from other cells, this was also significantly associated (P=0.00019) with the degree of enrichment (Figure S8g).

These data show that the set of PVLAB genes that are upregulated in TL are also significantly enriched for both GWAS genes and psychoactive drug targets, highlighting the importance of regional differences (and especially TL) for neuropsychiatric disorders genes and response to psychoactive drugs in PVALB.

### Regional gene expression differences in PVALB neurons are associated with gene dysregulation in neuropsychiatric disease

To assess if regional differences in gene expression in healthy brain can be relevant to disease-associated changes in brain tissue, we leveraged publicly available bulk RNA-seq data generated from neocortex tissue of diseased and control patients (Figure 5a), including BP, SCHI, ASD, OCD and major depressive disorder (MDD). All the DEGs identified between cases and controls are listed in Table S5. We first tested if the DEGs between TL and FL are enriched in the DEGs found in disease brain, and found the highest enrichment for neuronal cell types (Figure S9a). The most significant overlap with the DEGs identified in each brain diseases was detected for the PVALB sub-cluster, particularly for OCD (adjusted P = 2.7×10^−3^), BP (adjusted P = 9.6×10^−5^) and SCHI (adjusted P = 2.6 × 10^−5^) (Figure 5b). We also combined all disease related genes (non-redundant set of DEGs across all diseases, n=4,028), which confirmed the strongest enrichment for the DEGs in the PVALB sub-cluster. For the set of overlapping genes, we investigated the relationships between the expression changes in TL/FL (i.e., log_2_FC(TL/FL)) in PVALB and the changes in disease brain (i.e., log_2_FC (disease/control)). We observed that most of the PVALB genes that are up regulated in TL are also up regulated in BP (54.3%). In contrast, the majority of genes up regulated in TL showed downregulation in OCD (79.5%) and SCHI (66.8%) brain cortex (Figure 5c, d, e). Among the set of DEG in TL/FL and disease/control brain tissues, we identified several GWAS genes for neuropsychiatric disorders (tables in Figure 5c, d, e), with most of them being in common to at least two disorders. For example, we report *GRIN2A* and *HCN1* genes being both associated with schizophrenia by GWAS (P<10^−8^) and targets for psychoactive drugs (GRA and ARA), and consistently upregulated in TL and dysregulated (upregulated)in brain tissue from schizophrenia patients (bold font, tables in Figure 5c, d, e, f).

**Fig. 5.**
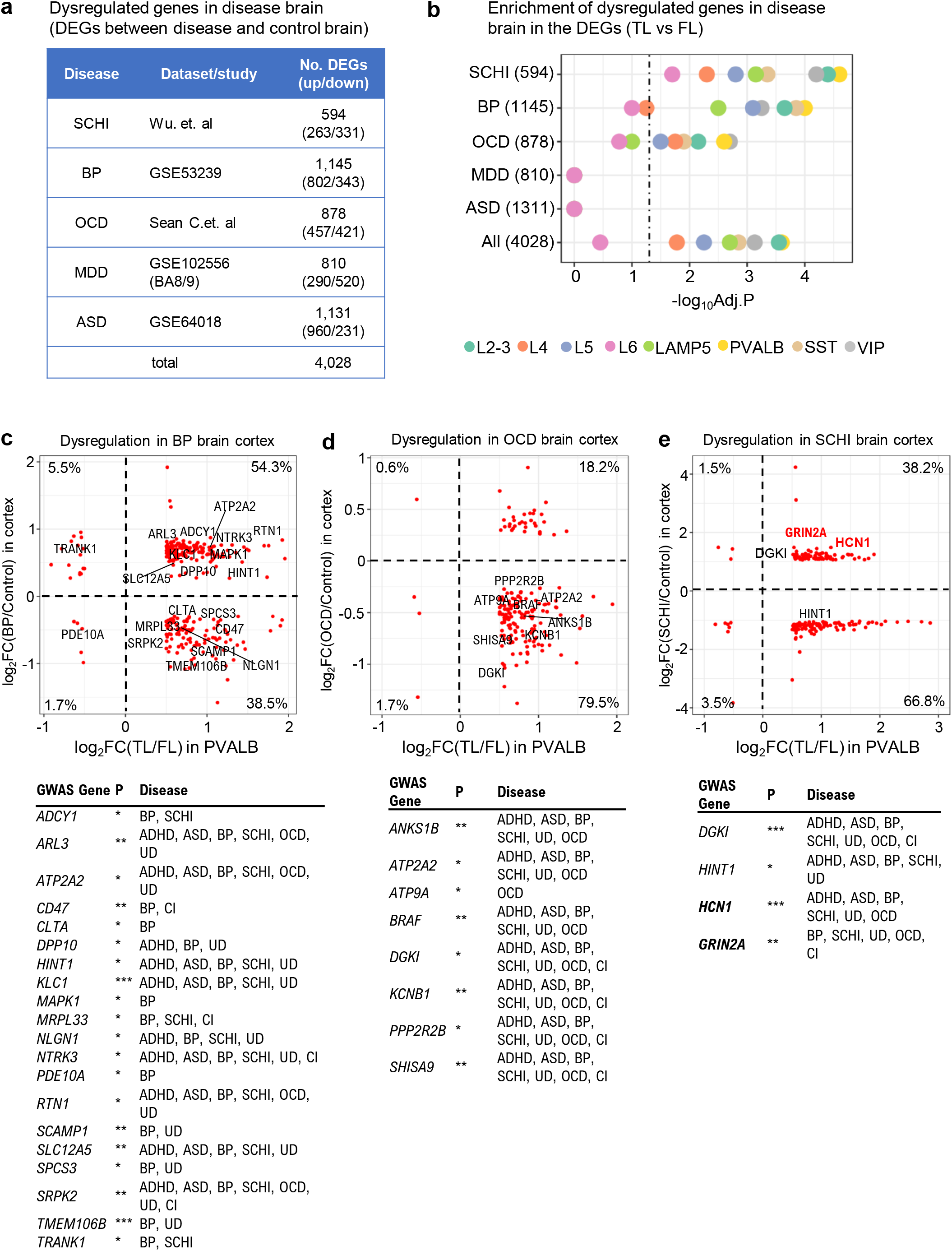
Genes expressed in PVALB neurons from the temporal lobe are also dysregulated in neuropsychiatric disease brain tissue. **a**. Details about the DEGs (absolute log_2_FC>0.25, FDR<0.05) identified in the brain of neuropsychiatric disease patients (see Methods for details). **b**. Bubble plot showing the enrichment of DEGs (absolute log_2_FC>0.5, FDR<0.05) between TL and FL (for each neuron subtypes) in the set of DEGs detected in the brain of neuropsychiatric disease patients Numbers in brackets are number of DEGs (disease vs control brain). Dotted vertical line indicates the threshold for statistical significance of enrichment (BH-adjusted P=0.05). **c, f**. Scatter dot plots showing the relationship between log_2_FC(TL/FL) of PVALB-expressed genes (x-axis) and log_2_FC (disease/control brain) (y-axis) for the set of genes significantly overlapping genes. Three scatter plots for BP (c), OCD (d) and SCHI (e). For each disease, the genes highlighted in each panel are GWAS-associated genes with GWAS P<10^−8^, and the numbers in each quadrant are the proportion of genes over the total number of DEGs in T/FL and in disease/control brain. Each Scatter dot plots is accompanied by a table showing the associated neuropsychiatric disease for each gene by GWAS: schizophrenia (SCHI), bipolar disorder (BP), cognitive impairment (CI), unipolar depression (UD), autism spectrum disorder (ASD), obsessive compulsive disorder (OCD), and attention deficit hyperactivity disorder (ADHD); for each gene we report the GWAS-association P-values: *, 10^−8^-10^−5^; **, 10^−8^-10^−10^; *** <10^−10^ ***.

## DISCUSSION

Psychiatric disorders are complex, poorly understood and difficult to diagnose. GWAS have confirmed the complex genetic etiology underlying these disorders, and identified manifold susceptibility genes. While substantial genetic heritability is shared across different psychiatric disorders^59^, the specific common genetic pathways are poorly defined. Moreover, the functional and cell type specific impact, also referred to as “vulnerability” ^18^, of these susceptibility genes is largely uncharted, and remains to be elucidated. In this study, we have generated a comprehensive single cell atlas of gene expression changes in TL and FL from fresh human brain samples. These single cell data have been integrated with (*i*) genetic susceptibility data for 7 psychiatric disorders (ADHD, BP, ASD, OCD, UD, SCHI, CI), (*ii*) RNA-seq data from healthy and disease human brain cortex (BP, SCHI, ASD, OCD, MDD) and (*iii*) drug-specific gene targets for common psychoactive drugs. Our integrative analysis allowed to determine cell type-specific vulnerabilities to disease genes and druggabilities to psychoactive drugs, which overall highlighted a specific neuronal subtype (PVALB) (Figure 6 – graphical abstract).

**Figure 6.**
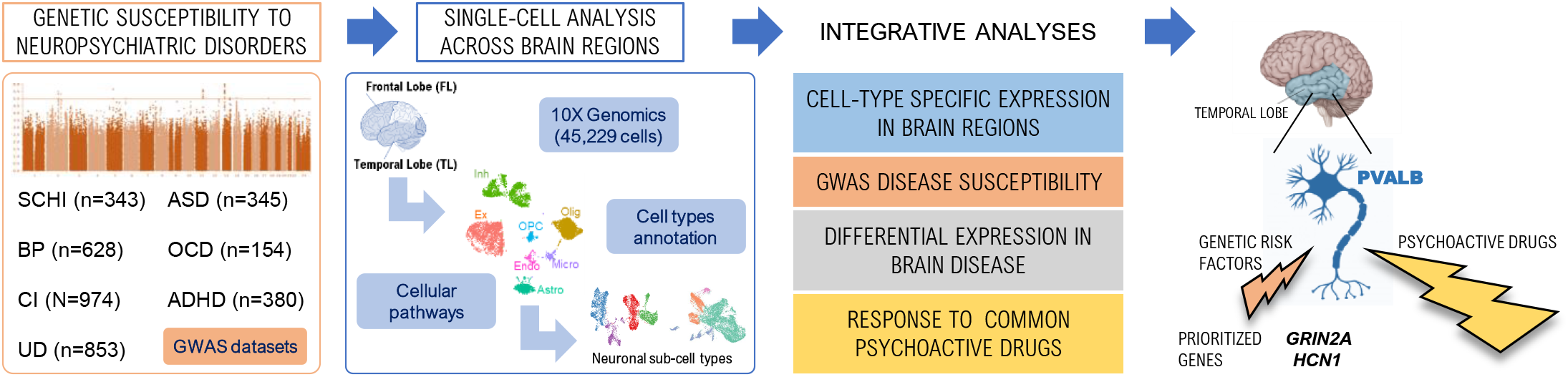
graphical abstract.

We started from defining the pathways shared between GWAS genes across different neuropsychiatric disorders, such as GABAergic synapse and dopaminergic synapse which modulate several processes relevant to neurodevelopmental disorders, including neuronal proliferation, migration, differentiation and connectivity^60^. Moreover, genetic dysregulation of pathways such as Rap1 and PPAR signaling in the PFC is sufficient to drive stress-relevant cognitive and synaptic phenotypes, and may contribute to inflammatory processes driving neurodegenerative diseases^61, 62^. A sub-set of 38 GWAS genes was common to all psychiatric disorders considered here; this includes genes such as *GRIN2A, DGKI*, and *SHISA9*, which are previously known to be involved in multiple psychosis^63, 64^. These findings suggest that these diverse psychiatric disorders have a genetic relationship with each other, and share genetic pathways and susceptibility genes, suggesting a common genetic basis of these diseases^65^.

We then postulated that these psychiatric disorders might also share a sub-set of common, more vulnerable brain cell types. By integrating our single cell with GWAS genes data, we found that neurons are the most vulnerable cell type to genetic susceptibility, particularly inhibitory neurons. Altered inhibitory control is believed to lead to a change in the power of gamma oscillations^66^, which is important in schizophrenia. Moreover, increase in the ratio between excitation and inhibition, called the E/I balance has been reported in autism spectrum disorder and children with autism show a reduced gamma frequency modulation to a visual task, whereas increased synchrony was observed in ADHD^52, 67^.

Amongst different neuronal subtypes, we found PVALB, SST and L5 neurons having the highest enrichment for GWAS genes, suggesting that disruption the specific gene biological process in these cells may contribute to neuropsychiatric disorders. These neuronal sub-types and the biological processes they modulate have been previously associated with many psychiatric diseases. PVALB cells are believed to activate pyramidal neurons only if the signal from excitatory neurons is sufficient and optimize the signaling in both EX and INH^68^. SST neurons gate excitatory input onto pyramidal neurons within cortical microcircuits, mainly coming from L5 layer of excitatory neuron^69^. These signaling processes, when dysregulated, have been implicated in psychiatric diseases^70^.

Regional differences between FL and TL are important for psychiatric disorders. The FL is responsible for shaping social behavioral characteristics such as personality, decision-making, motivation, while the TL is typically related to emotional regulation, planning, reasoning and problem solving^71^. Here we annotated regional differences in gene expression at the single cell level, and tested if these might contribute to explain psychiatric disease susceptibility by GWAS genes. First, our study points to important regional gene expression differences (FL *vs* TL) in functional pathways affecting both neuronal and sub-neuronal cell populations. Second, we show GWAS genes for psychiatric diseases being enriched in PVALB cells from the TL, which suggests that shared co-morbidities related to memory deficits might be due, at least in part, to functional dysregulation of PVALB neurons. Memory disorders have been primarily related to destruction in PVALB^72^, and defective signal transmission in PVALB neurons has been associated with higher likelihood of developing cognitive deficits^73^.

More generally, transcriptional changes between FL and TL were related to multiple signaling pathways (e.g., cGMP−PKG, cAMP, long term potential) and pointed to widespread structural plasticity (of axons, dendrites and synapses). For example, we identified regional gene expression differences impacting synaptic vesicle cycle genes specifically in PVALB neurons (including SNAP-25 and VAMP-2, members of the SNARE complex involved in endocytosis of synaptic vesicle, associate to ADHD)^74^. Our findings are also in keeping with recent work showing a temporal specificity of synaptic change in schizophrenia, with synaptogenesis being predominant earlier in the disease, and synaptic loss in chronic phases^75^. Beyond ADHD and BP, a decrease of PVALB inflammatory neurons has been reported in bipolar patients, and this may alter intrinsic inhibitory networks within the superficial layers of the TL, which might disrupt integration and transfer of information from the cerebral cortex to the hippocampus^76^.

Since the regional gene expression differences were identified in brain tissue from patients without history of psychiatric disease, we investigated to which extent these intrinsic changes correlate with gene expression differences in disease brain or in response to psychoactive drugs. Enrichment for genes dysregulated in disease brain and for psychoactive drug gene targets was the strongest for the inhibitory neuronal sub-type, in most cases with specificity to the neurons from TL. TCA and SSRI showed high and similar enrichment in PVALB from TL, which is not unexpected given that TCA and SSRI have similar structure and efficiency^77^. Activity of vesicular glutamate transporters (VGLUTs), responsible for the uptake of glutamate into the synaptic vesicle^78^, is increased in TL in schizophrenia and bipolar patients after treatment with SSRI^79^. Our identification of PVALB neurons as strongly enriched for genes dysregulated in brain from schizophrenia patients is also in keeping with recent studies in adult Macaque cortex sowing schizophrenia DEGs in excitatory neuron subtypes, where the PVALB-type was the most affected inhibitory subtype^80^.

Our integrative analyses identified two key genes, *GRIN2A* (coding subunit 2A of the NMDA-type receptor (NMDAR) and target of Glutamate Receptor Agonists (GRA)) and *HCN1* (coding the channel mediating the h-current (Ih) and target of Adrenergic Receptor Agonist (ARA)), as being upregulated in PVLAB neurons from TL, upregulated in brain cortex from schizophrenia patients, and strongly associated with schizophrenia (and other psychiatric disorders) by GWAS (10^−10^< GWAS P <10^−8^). Schizophrenic psychosis may originate in the TL of due to the deficits in establishment of synaptic contacts which is caused by PVALB dysfunction, which is mediated by *GRIN2A* in PVALB-positive interneurons^81^, and has been proposed for therapeutic targeting. Neural oscillations are abnormal in schizophrenia patients compared with healthy controls, and *HCN1*, a hyperpolarization-activated cation channel that contributes to the native pacemaker currents in neurons, has been associated with schizophrenia^82^, schizophrenia endophenotypes^83^ and working memory in schizophrenia patients^84^. Therefore, by integrating regional gene expression differences at single cell level with GWAS data, gene dysregulation in disease and response to psychoactive drugs we highlighted *GRIN2A* and *HCN1* function in PVALB neurons in the TL.

While our analyses allowed us to define disease-relevant cell types for psychiatric disorders genes, there are limitations to our approach. First, our study stemmed from the single cell analysis of only 3 bio samples, which will need to be replicated in larger patient cohorts to better resolve transcriptomic changes in TL and FL. However, our analysis yielded >45,000 cells and high sequencing depth (median UMI per cell = 825, UMI of 1/3 cells is over 1000)^85^, providing high quality single cell data into which the other data (GWAS, bulk-RNA seq, psychoactive drug targets) are integrated. Second, we only included GWAS data for 7 psychiatric disorders, and we used a relaxed threshold for the GWAS significance of the genes (GWAS P-value < 10^−5^). However, this was used only at the first stage of the analysis to collect a large set of GWAS genes for integrative analyses. The GWAS genes were then filtered (using other datasets) and further refined using more stringent thresholds (GWAS P-value < 10^−8^) at later stage of our analysis. Third, as the single cell data were acquired by nuclei sequencing, this might have limited the analysis as RNA species from other cellular compartments were not captured.

In summary, we generated a single cell-resolved dataset from temporal and frontal lobe tissues, which provided new insights into the regional brain differences and cell type-specific vulnerabilities to neuropsychiatric disorders and drugs. Recent studies investigated the cell type vulnerability to psychoactive drugs^18^, highlighting the role or SST and VIP interneurons. Our study identified additional sets of neurons (PVALB and LAMP5) as highly vulnerable cells to psychoactive drugs and show this vulnerability is specific to neurons from the TL. Integrating brain single cell with GWAS data, Olislagers *et al*.^86^ showed that neuronal cell subsets are consistently implicated in several psychiatric disorders. Our study confirms and extend these results, as we further refined the neuronal subtypes to the inhibitory sub-type (PVALB and SST) and again highlight their specificity to the TL. However, differently from both ^18^ and ^86^, in our study we jointly integrated multiple datasets: regional single cell analysis in TL/FL, GWAS data, gene expression (bulk-RNA) in brain from psychiatric disorders patients and gene response to psychoactive drugs. This allowed us to disentangle the relative contribution of genetic susceptibility and psychoactive drugs the cell type vulnerability. Overall, of the genes upregulated in PVALB from TL, we found 9.8% being enriched for GWAS genes and 35.6% being enriched for psychoactive drug targets, respectively. This suggests a considerably larger contribution (3.6-fold) of psychoactive drugs to PVALB cell vulnerability in the TL. Notably, neither GWAS or psychoactive drug target genes were enriched in the PVALB cell population from the FL, highlighting the functional specialization of the two lobes. Finally, we provide a unique data resource and present a detailed integrative single cell analysis of cellular heterogeneity and brain regional changes of psychiatric disease genes, which will enable specific investigations of disease risk genes and their cell type-specific contribution to disease susceptibility.

## ACKNOWLEDGEMENTS

This study was funded by Scientific Research Foundation for Advanced Talents (No. 3012000223), the National Natural Science Foundation of China (No. 81972153, 81901955, 82120108018), Natural Science Foundation of Jiangsu Province (No. BK20191066), Priority Academic Program Development of Jiangsu Higher Education Institutions (No, JX10231804), and Gusu School Nanjing Medical University (GSKY202201010), National Key Research and Development Program of China (No.2021YFA1101802).

