## Supplementary Figures 1-10 for "Decoding frontotemporal and cell type-specific vulnerabilities to neuropsychiatric disorders and psychoactive drugs"

### Figure S1

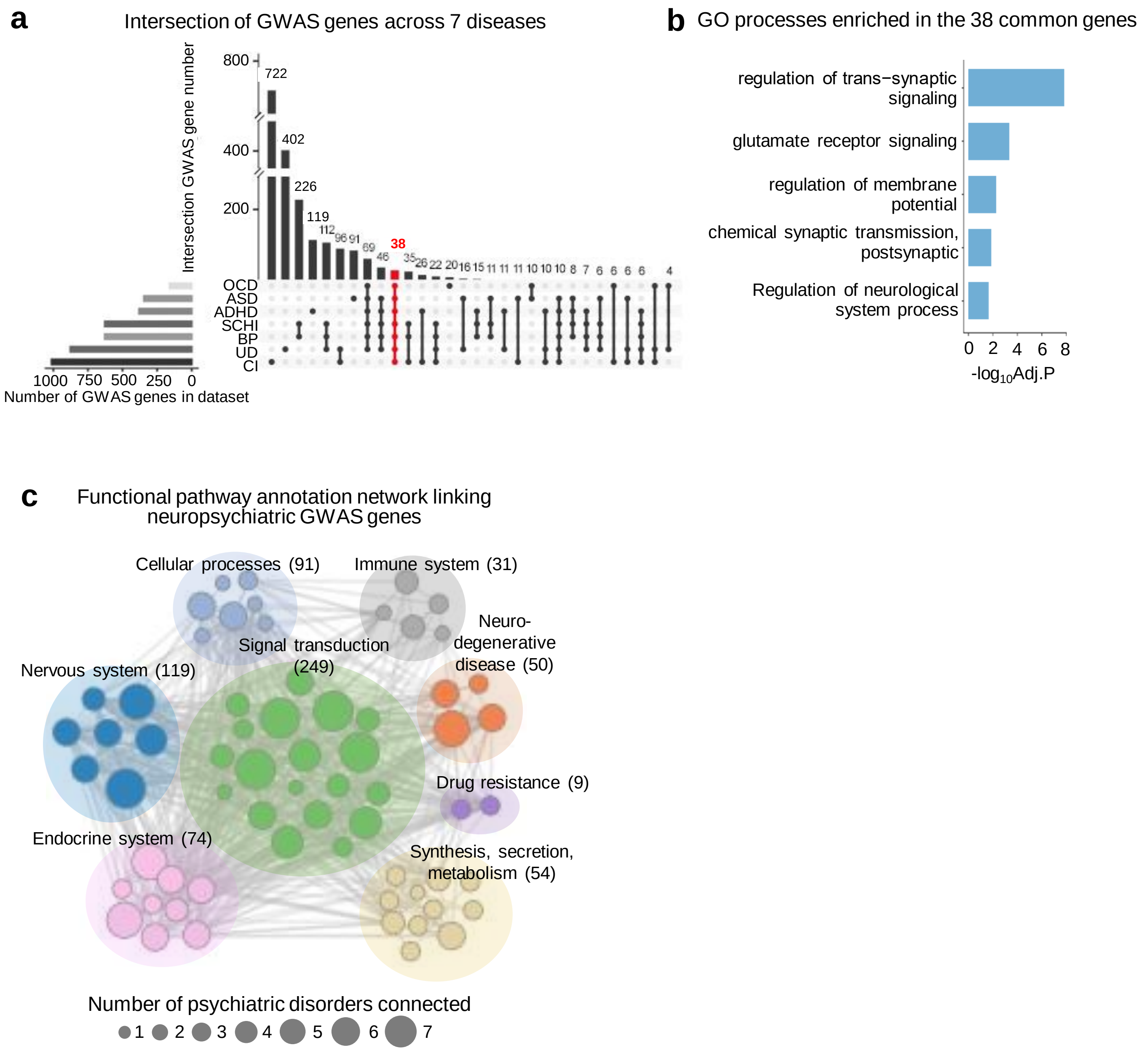


**Figure S1. Functional annotation of GWAS genes for neuropsychiatric disorders identifies common pathways and 38 common GWAS genes. a.** Upset plot is an alternative venn plot to show the integrated GWAS genes across 7 diseases considered here, 38 genes (highlighted in red) were shared by all diseases. **b.** Non-redundant Gene Ontology (GO) processes that are significantly enriched (BH-adjusted P<0.05) in the set of 38 common GWAS genes. **c.** KEGG pathways enriched (BH-adjusted P<0.05) in the GWAS genes that are classified to 8 main groups according to the KEGG database. Dots size represents the number of psychiatric diseases with significant enrichment for the same pathway. Detailed list of enriched pathways is in Table S2.

### Figure S2

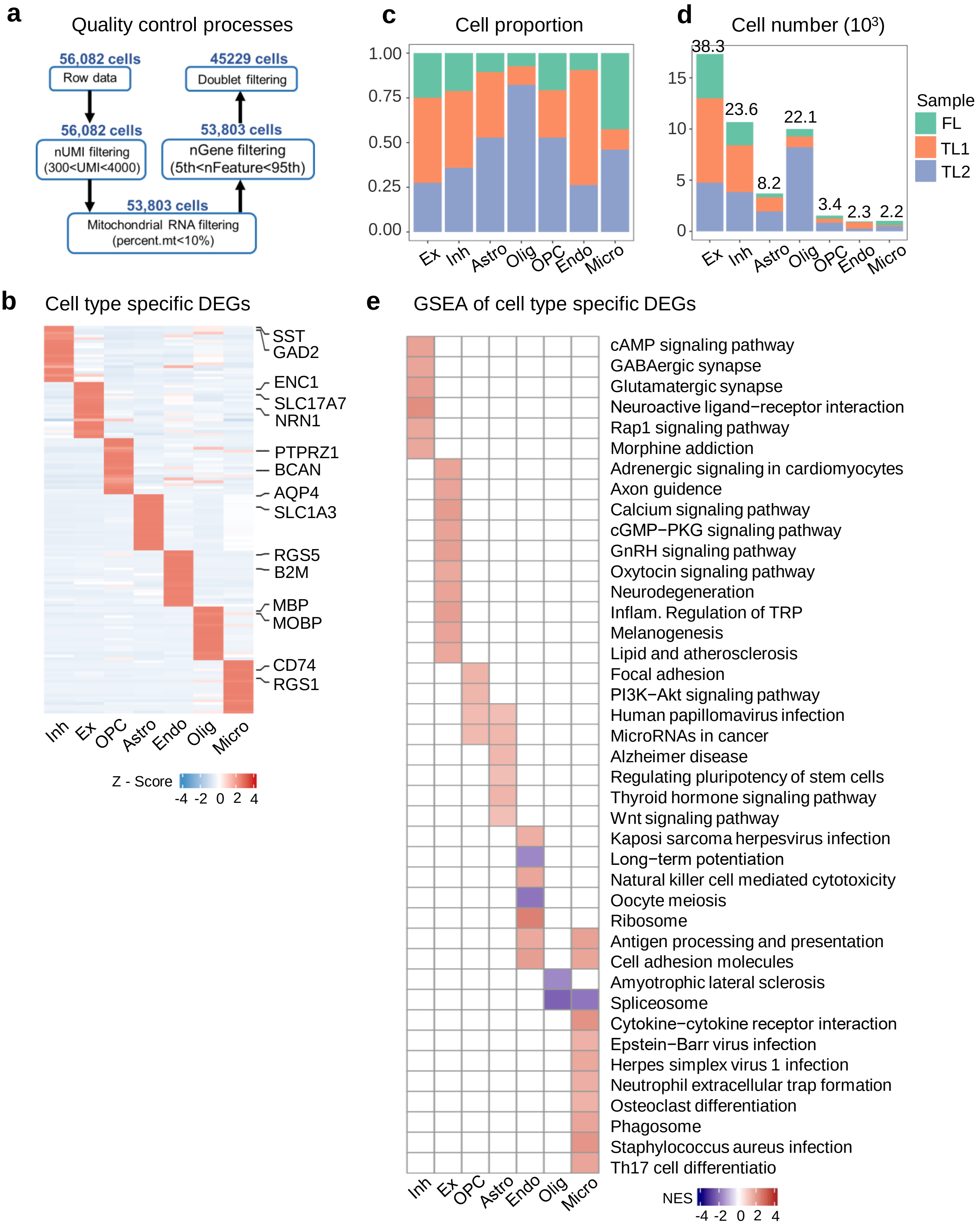


**Figure S2. Description of snRNA-seq data. a.** Flow chart showing the quality control (QC) processes used to analyze the snRNA-seq data (see Methods for additional details). **b.** Single cell level heat map exhibiting cell type-specific gene marker expression (z-scored) in each cell clusters (average $log_2FC>0.5$, BH-adjusted $P<0.05$). **c.** Proportion of cells belonging to each sample in each cell type. **d.** Number of cells belonging to each sample in each cell type with relative proportion over all cells. **e.** Gene set enrichment analysis (GSEA) of cell type-specific differentially expressed genes (DEGs) across all cell types colored by degree of enrichment, i.e. normalized enrichment score (NES) (BH-adjusted $P<0.05$).

### Supplementary Figure 3

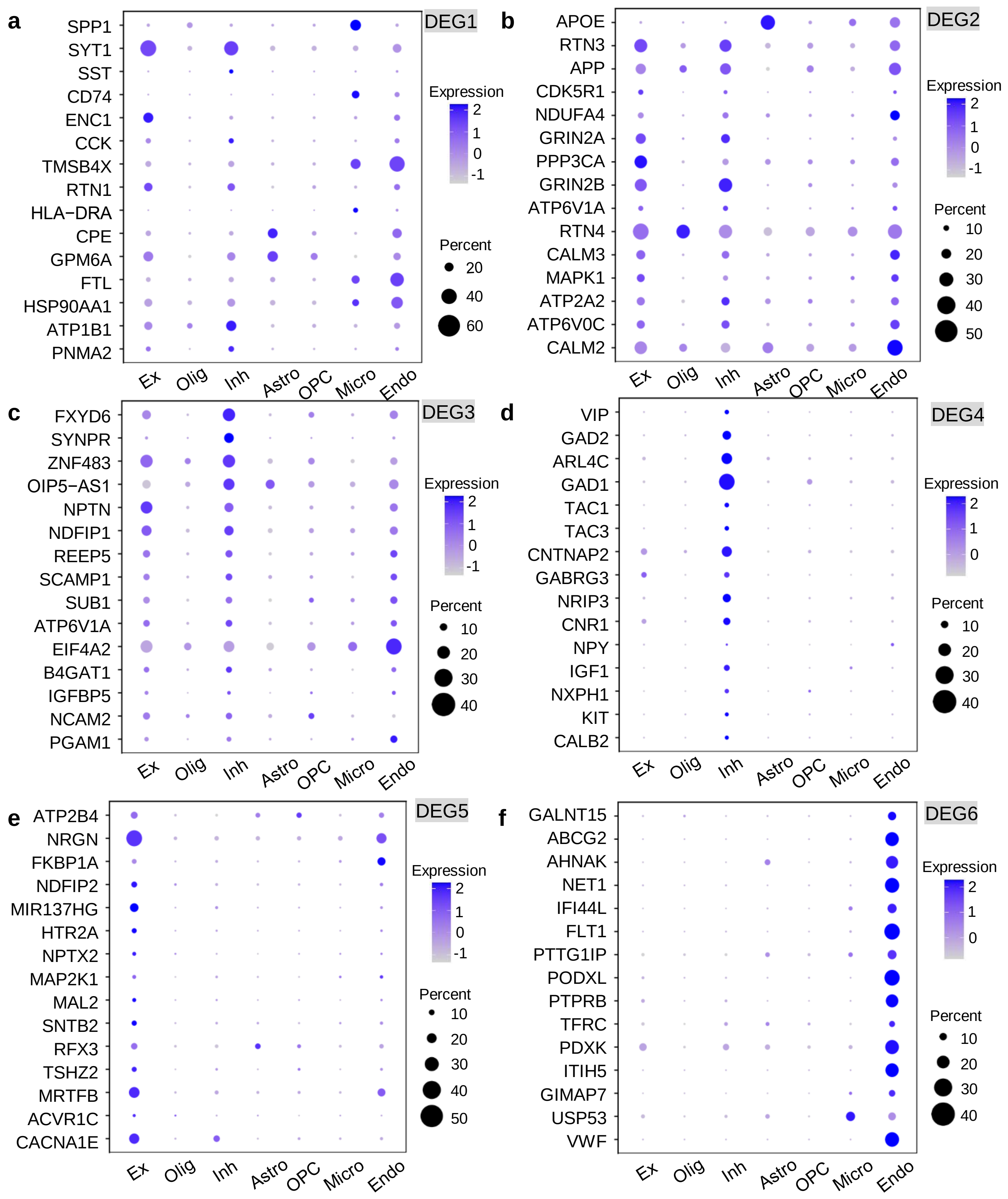


**Figure S3. Expression level of regional DEGs in each cell types. a-f.** Bubble plot showing the top 15 DEGs between FL and TL within the DEG1-DEG6 clusters (reported in Figure 2c), where the bubble color represents the gene expression level, and the bubble size is the proportion of cells expressing the gene. The DEG4 cluster, including glutamate receptors (*GAD1*, *GAD2*) and GABA receptor (*GABRG3*), is specific to INH, while the DEG5 cluster, including neurexin (*NRGN*), serotonin receptor (*HTR2A*) and calcium signaling factors (*NPTX2*, *CACNA1E*), is specific to EX (Figure 2c and Figure S3d, e). In contrast, the DEG2 cluster was found in multiple cell types, and include INH- and EXH-expressed genes such as *GRIN2A*, *GRIN2B*, and other genes such as *APOE*, *APP*, and *MAPK1* (Figure 2c and Figure S3b). The DEG3 cluster was composed of genes mostly expressed by EX and INH, whose function is enriched for dopaminergic synapse and synaptic vesicle cycle processes (Figure 2c and Figure S3c). Other clusters (e.g., DEG6) were expressed in Endo from FL, and are involved in immune response processes (Figure 2c, Figure S3f).

### Figure S4

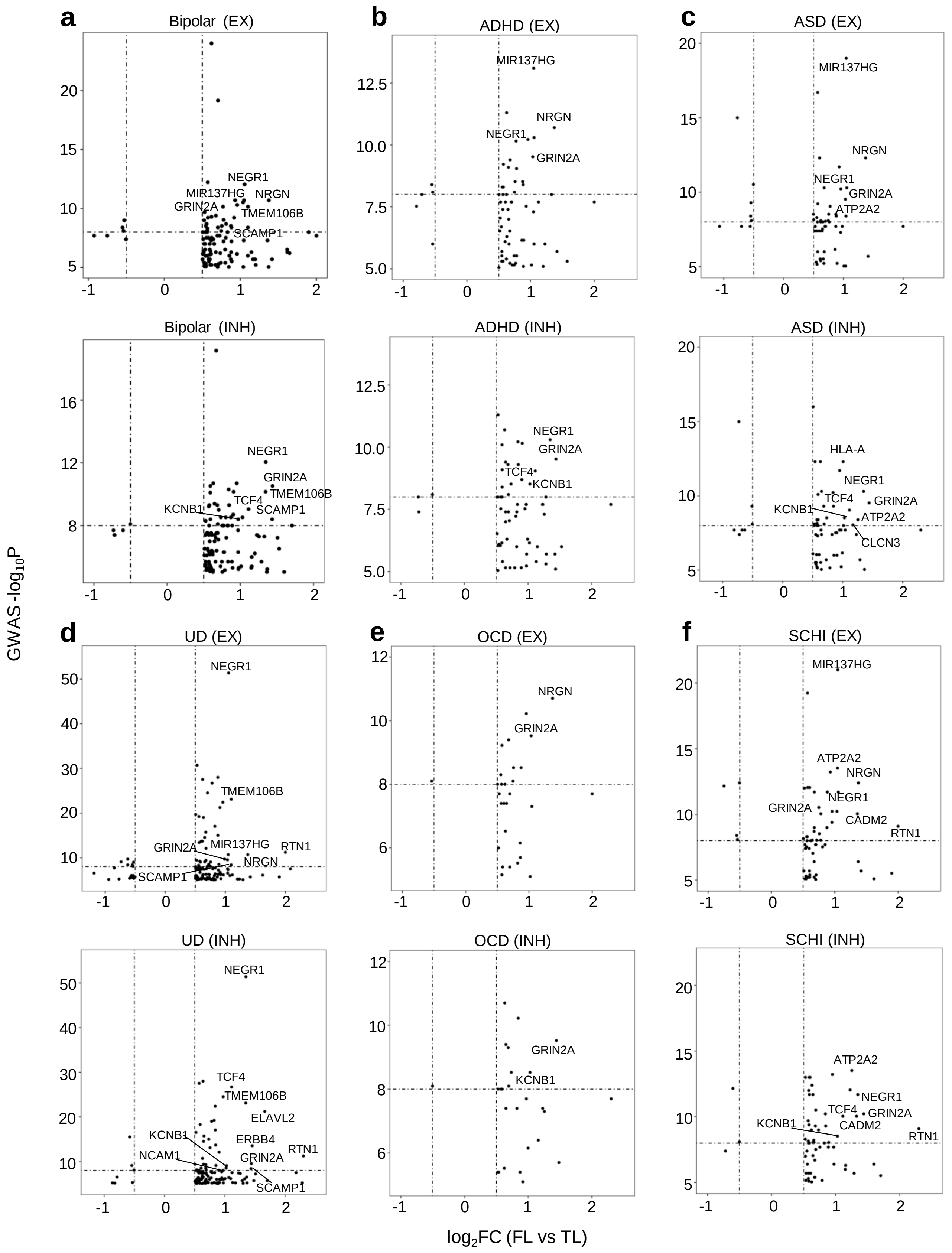
a
Bipolar (EX)
Bipolar (INH)
b
ADHD (EX)
ADHD (INH)
c
ASD (EX)
ASD (INH)
d
UD (EX)
UD (INH)
e
OCD (EX)
OCD (INH)
f
SCHI (EX)
SCHI (INH)
NEGR1
MIR137HG
NRGN
GRIN2A
TMEM106B
SCAMP1
KCNB1
TCF4
HLA-A
ATP2A2
CLCN3
CADM2
RTN1
ELAVL2
ERBB4
NCAM1
GWAS -log10P
log2FC (FL vs TL)

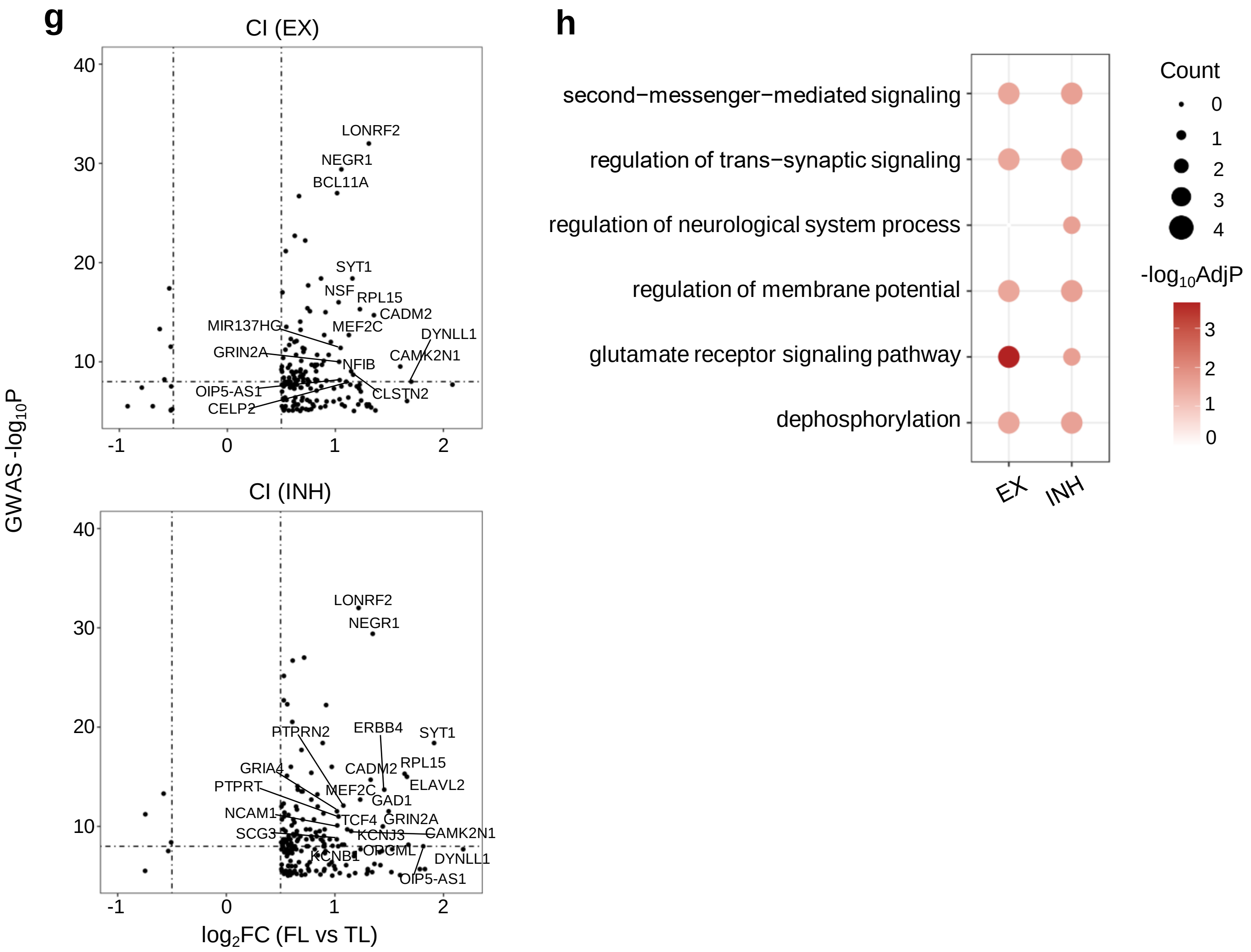


**Figure S4. Relationship between genes $\log_2$FC (FL and TL) and significance of GWAS associations in neuronal cells. a-g**. Scatter dot plot showing the relationship between $\log_2$FC (FL and TL) and GWAS P-values for genes associated with BP, ADHD, ASD, UD, OCD, SCHI, and CI in EX (top) and INH (bottom) neurons, respectively. Strongly associated genes with $\log_2$FC>1 and GWAS $P<10^{-8}$ in each disease are highlighted. **h**. GO biological processes significantly enriched (BH-adjusted P<0.05) in the differentially expressed (TL/FL) set of 38 common GWAS genes.

### Figure S5

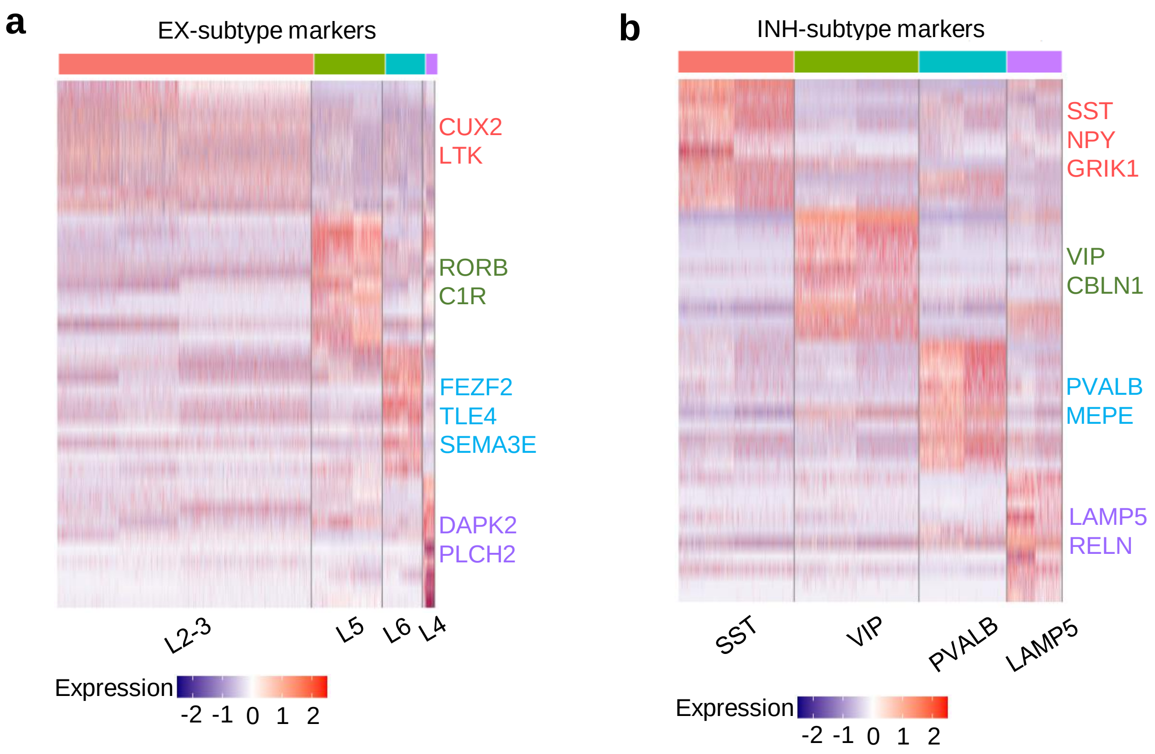


**Figure S5. Annotation of excitatory and inhibitory neuronal subtypes. a, b**. Heatmap showing the normalized gene expression of subcluster-specific genes (average $log_2FC>0.5$, BH-adjusted $P<0.05$) in EX (a) and INH (b) cells.

### Figure S6

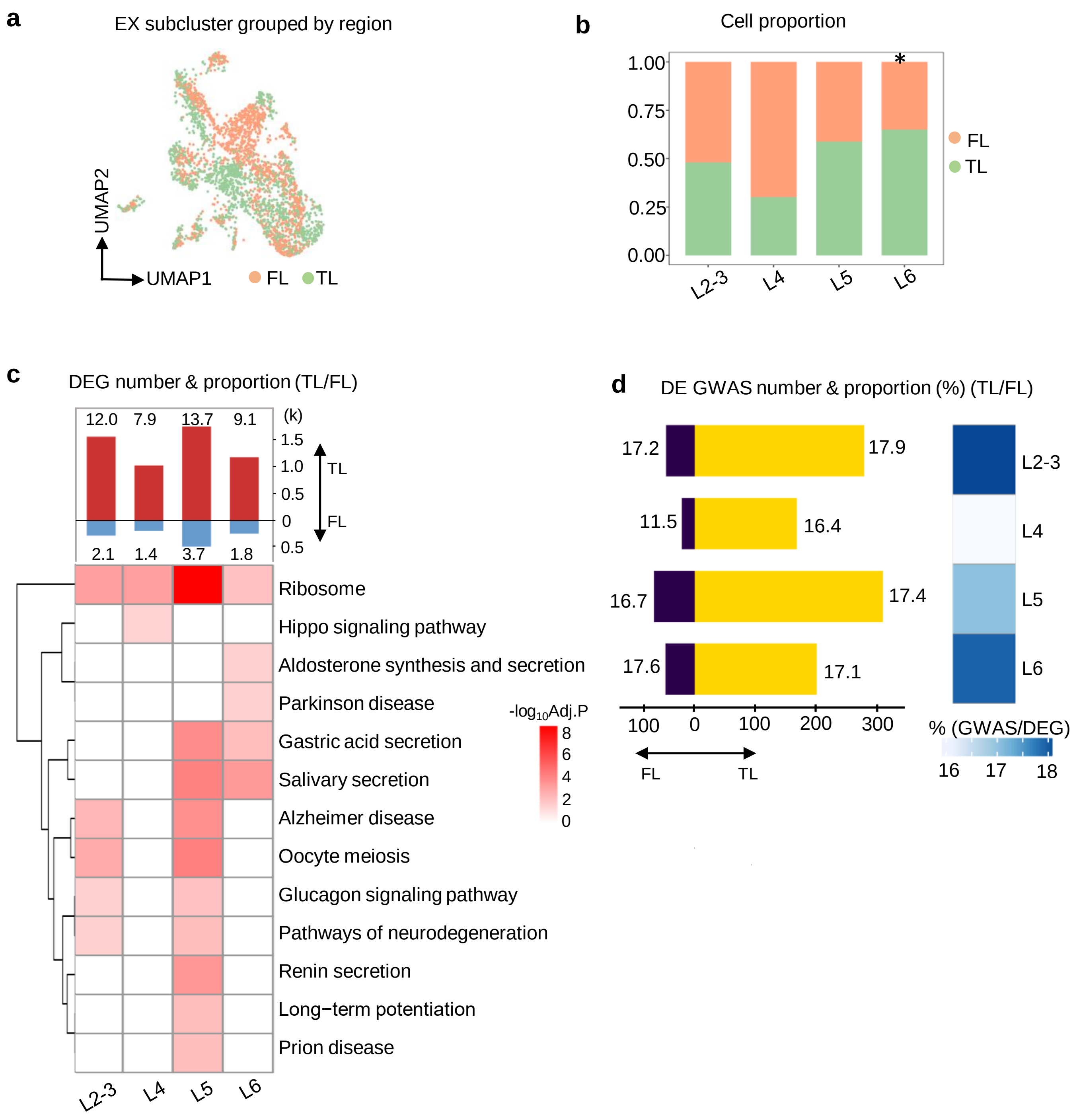


**Figure S6. Transcriptional changes in EX neuronal subcluster between FL and TL, and enrichment for GWAS genes. a.** UMAP plot showing the two regions merge completely after doing batch correction. **b.** Cell proportion of FL and TL in each subcluster and the significant cluster by chi-square test are labelled with asterisks ($P<0.05$). **c.** Number of DEGs between FL and TL, where the ratio (%) of DEGs over the total number of genes expressed in each subcluster is reported (*top*). Heatmap exhibit the significantly enriched (BH-adjusted $P<0.05$) KEGG pathways in the set of DEGs between TL and FL. **d.** Number of GWAS genes that are DE between FL and TL (BH-adjusted $P<0.05$), with the proportion of DE GWAS genes over the total number of DEGs in each cluster (*left*). Proportion of all DE GWAS genes over total DEGs in each cell type (*right*). No significant enrichment for GWAS genes was found on the subsets of EX neurons.

### Figure S7

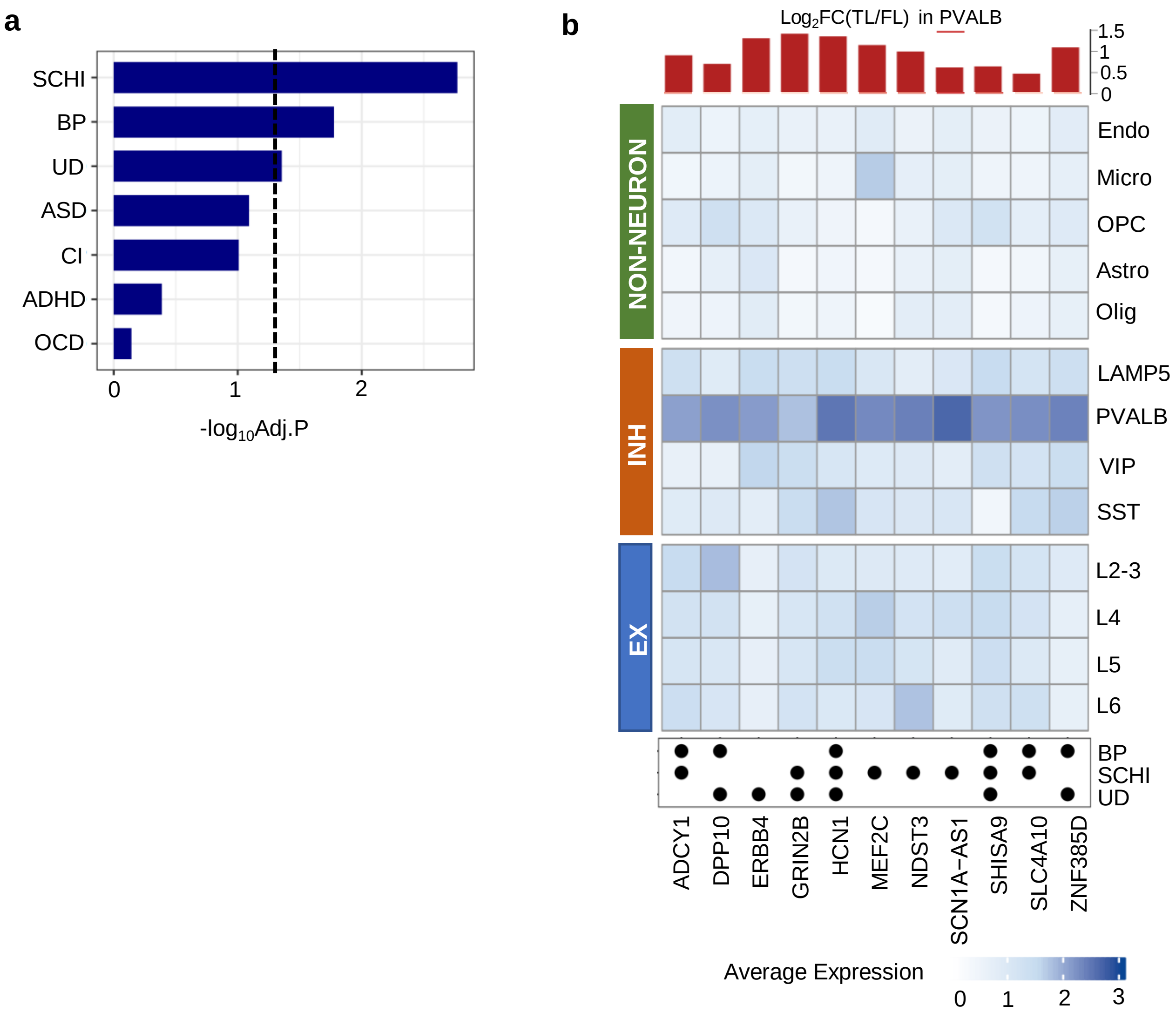


**Figure S7. Genes in PVLAB and upregulated in TL are enriched for GWAS genes related to schizophrenia and bipolar disorder. a.** Enrichment of GWAS genes for each neuropsychiatric disorder in the genes expressed in PVALB subcluster and significantly upregulated in TL vs FL (BH-adjusted P<0.05), with a dot line represents the significance threshold (hypergeometric test, BH-adjusted P<0.05). **b.** heatmap showing the average expression of GWAS-associated genes in UD, BP, and SCHI that are specific to PVALB cells and upregulated in TL across all cell types including INH subtype, EX subtype and non-neuronal cell types. Bar plot on the top reflect the gene fold change between TL and FL in PVALB neurons. Dot plot on the bottom reflects which disorder each GWAS gene belong to.

### Supplementary Figure 8

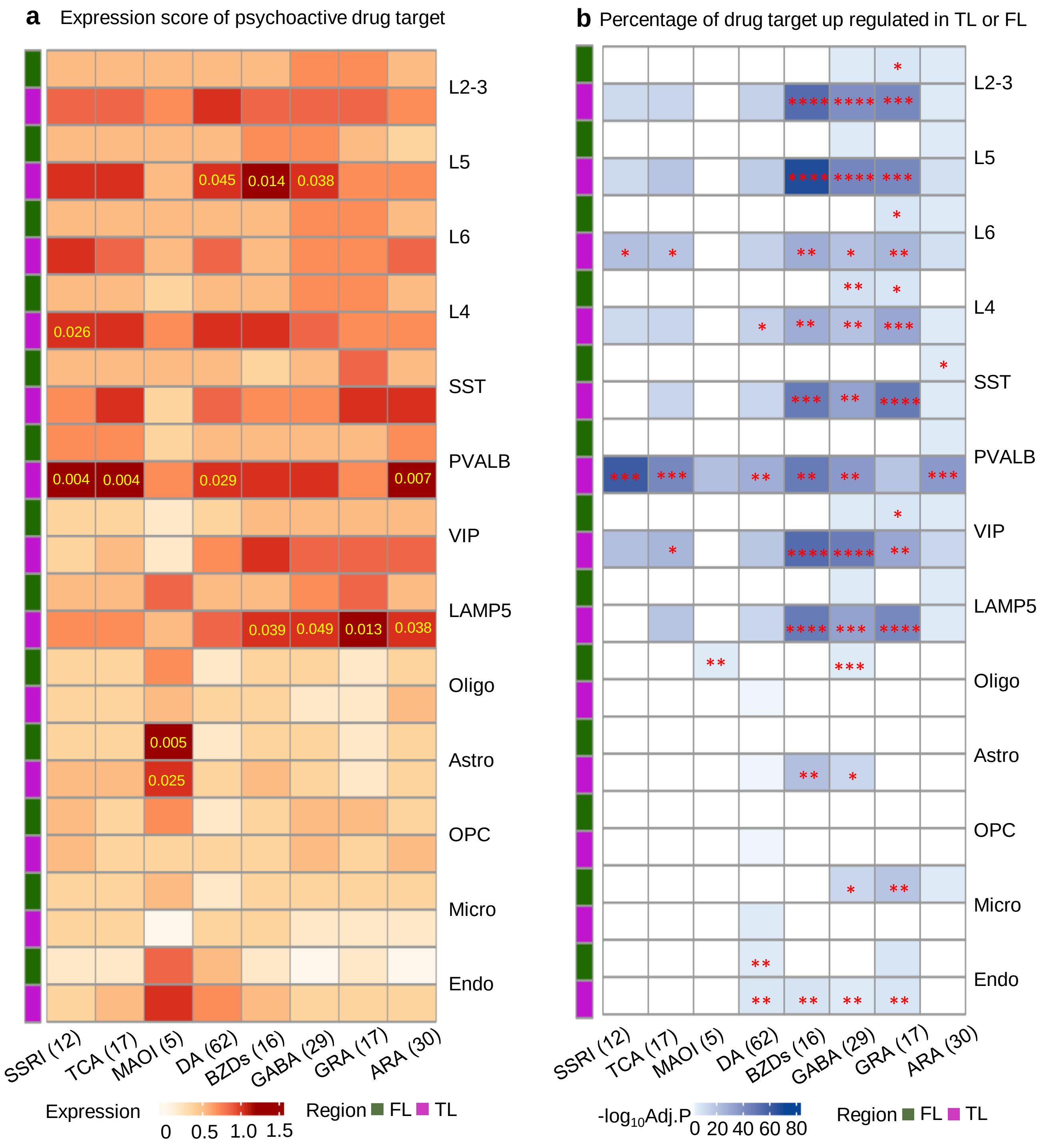


**Figure S8. Enrichment for psychoactive drug target genes reflecting cell vulnerability to psychoactive drugs. a.** Cell type resolved gene expression level of drug-target genes for each psychoactive drug group (class); numbers represent the number of target genes. Drug class names: SSRI: Selective Serotonin Reuptake Inhibitor, DA: Dopamine Receptor Agonists, TCA: Tricyclic Antidepressant, BZDs: Benzodiazepines, MAOI: Monoamine Oxidase Inhibitor, GRA: Glutamate Receptor Agonists, GABA: GABA Receptor Agonists, ARA: Adrenergic receptor agonist. The significant (BH-adjusted P<0.05) z-scores of enrichment for expression in a given cell type are indicates and the actual BH-corrected P-value is reported in the graph. **b.** Heatmap showing the percentage of drug target genes that are up regulated in TL or FL, respectively, across all cell types (including neuronal sub cell types). Significance of the enrichment (hypergeometric test) for differentially expressed drug target genes (between TL and FL) within each drug target group is indicated. Significance (P-value) thresholds: * 0.01-0.05, ** 0.001-0.01, *** 0.0001-0.001, **** <0.0001. **c-o.** Relationship between normalized expression level of drug-target genes (x-axis) and enrichment of drug-target gene overrepresentation (y-axis) in each cell type. P-values of significance of the association (correlation) between expression and enrichment of psychoactive drug-target gene overrepresentation are also reported. The most significant association (P=0.0002) is found for the psychoactive drug target genes in PVALB neurons from the TL. (c. L2-3, d. L4, e. L5, f. L6, g. PVALB, h. SST, i. VIP, j. LAMP5, k. Astro, l. Endo, m. Micro, n. Olig, o. OPC).

### Figure S8

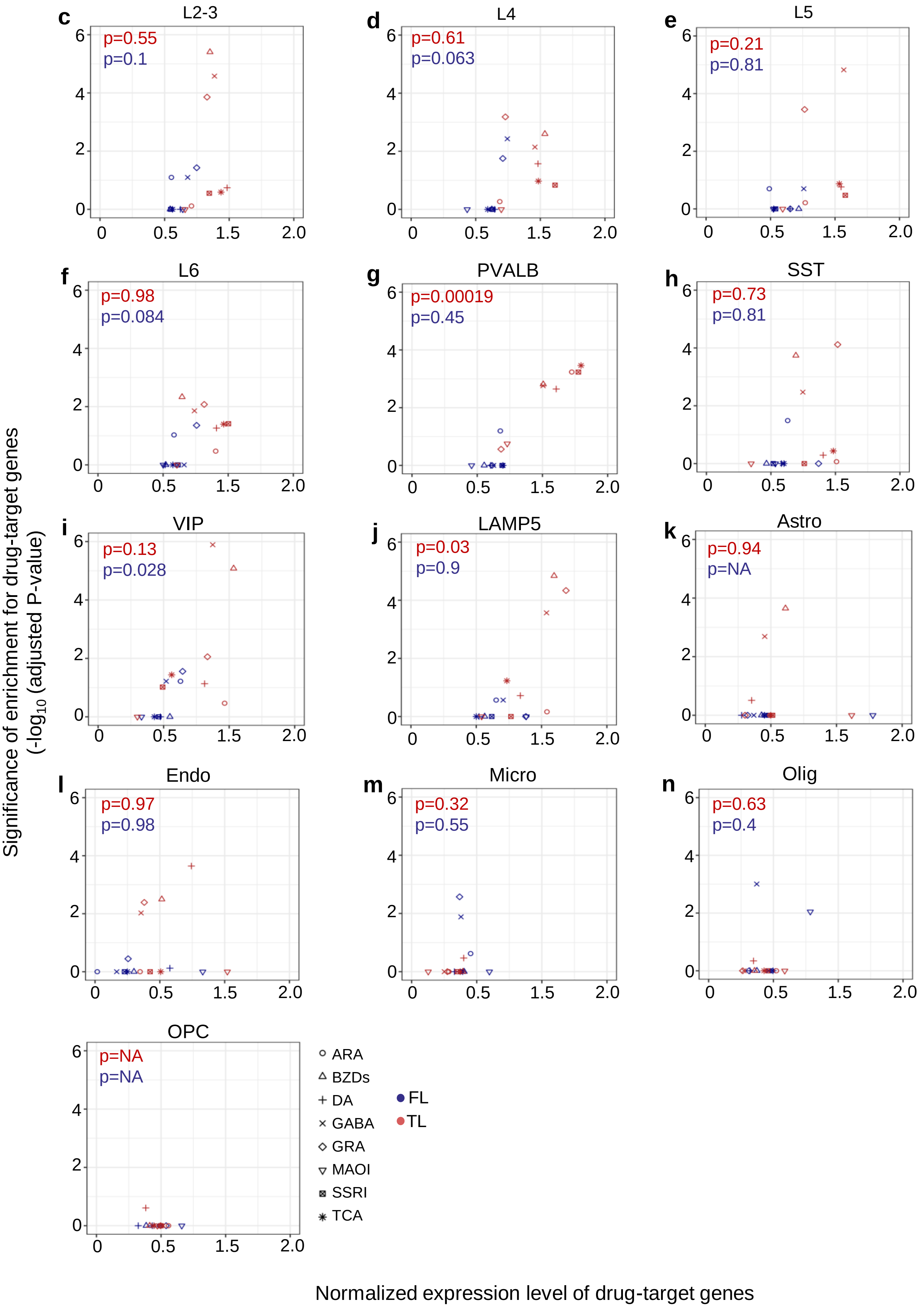
c
L2-3
p=0.55
p=0.1
d
L4
p=0.61
p=0.063
e
L5
p=0.21
p=0.81
f
L6
p=0.98
p=0.084
g
PVALB
p=0.00019
p=0.45
h
SST
p=0.73
p=0.81
i
VIP
p=0.13
p=0.028
j
LAMP5
p=0.03
p=0.9
k
Astro
p=0.94
p=NA
l
Endo
p=0.97
p=0.98
m
Micro
p=0.32
p=0.55
n
Olig
p=0.63
p=0.4
OPC
p=NA
p=NA
ARA
BZDs
DA
GABA
GRA
MAOI
SSRI
TCA
FL
TL
Significance of enrichment for drug-target genes
(-log10 (adjusted P-value)
Normalized expression level of drug-target genes

### Supplementary Figure 9

a

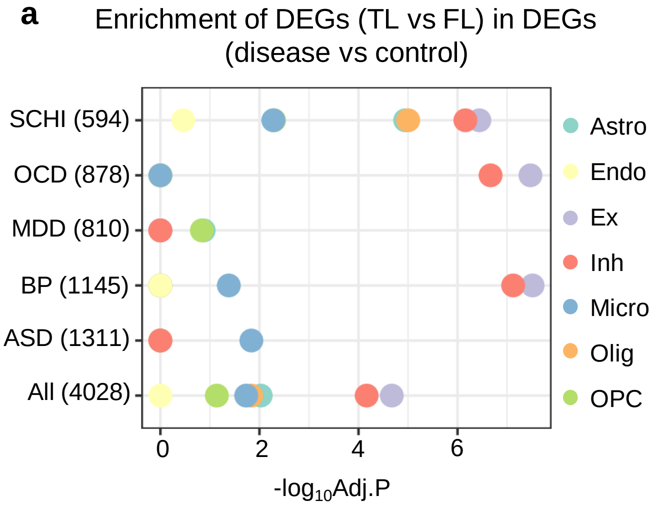


**Figure S9. Overlap of genes dysregulated in diseased brain and genes differentially expressed between TL and FL in each cell type. a.** Bubble plot showing the enrichment (x-axis, BH-adjusted P-value) of DEGs between TL and FL (absolute $log_2FC>0.5$ and FDR<0.05) in the set of dysregulated genes in disease brain tissue (DEGs between disease and control brain, absolute $log_2FC>0.25$ and FDR<0.05), retrieved from public dataset sand studies (see Methods for details). Numbers in brackets are number of DEGs (disease vs control) for each disease. The most significant enrichments are found in the neuronal cells. OCD showed no significant enrichments.

### Supplementary Figure 10

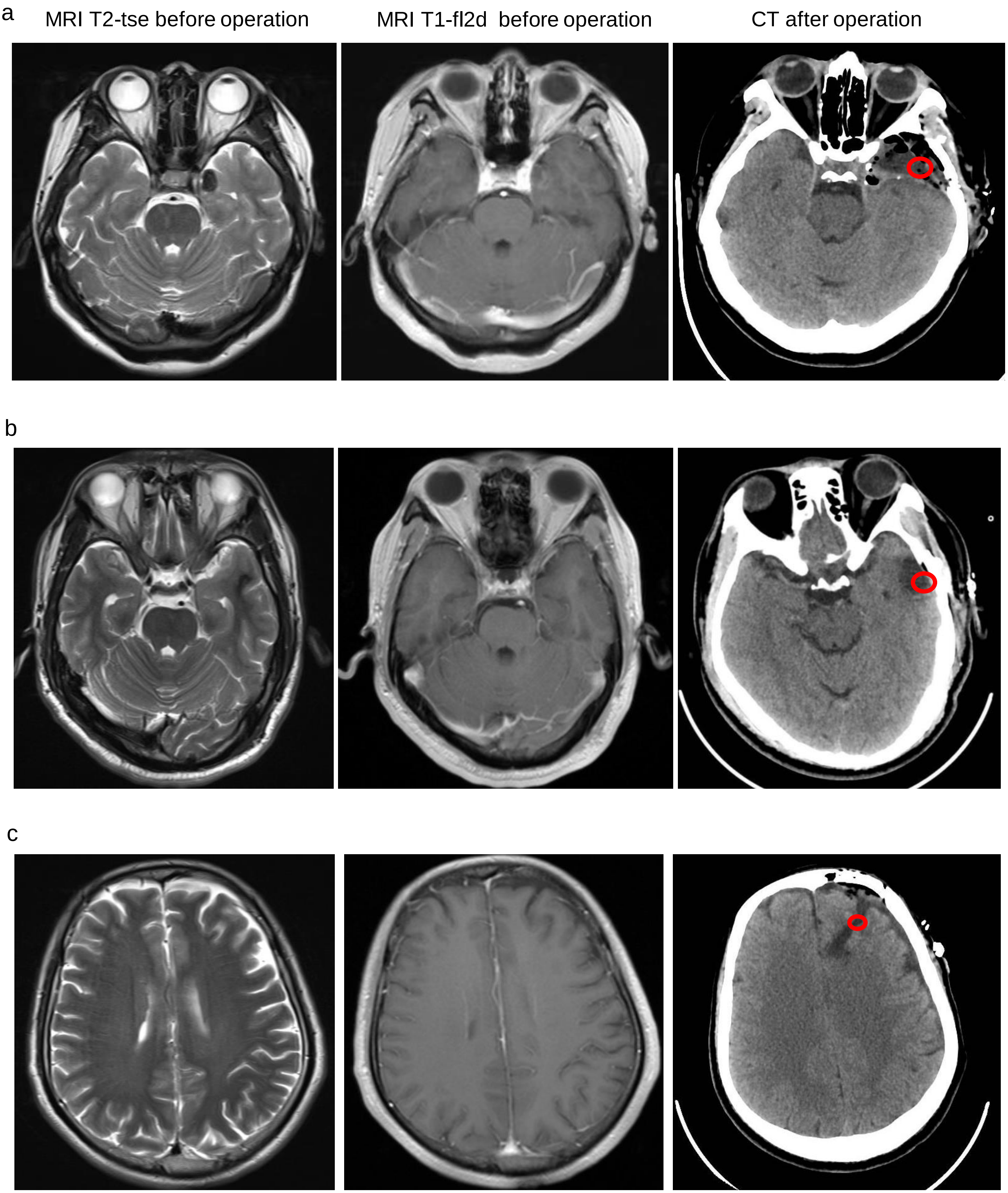


**Figure S10. Donors' brain MRI and CT images. a, b** CT images from TL1 and TL2 patient sample, **c** is from FL patient sample, containing preoperative MRI T2-tse, preoperative MRI T1-fl2d and postoperative CT images. The red circle is the sampling location for single-cell sequencing. More details on the patients in Table S3.
